# PIANO: Probabilistic Inference Autoencoder Networks for multi-Omics enables robust generative modeling of gene expression and scales single-cell integration to 100 million cells

**DOI:** 10.64898/2026.08.06.743394

**Authors:** Ning Wang, Christopher Cardenas, Victor E. Nieto-Caballero, David Turner, Hannah Feinberg, Dan Yuan, Nathaniel Scott, Michael DeBerardine, Shu Dan, Lakme Caceres, Jessica Schembri, Zizhen Yao, Changkyu Lee, Jonathan W. Pillow, Fenna M. Krienen

## Abstract

Single-cell RNA technologies enable the routine acquisition of transcriptomic atlases. However, these molecular profiles are influenced by overlapping sources of variation. Since these covariates confound comparisons, data integration is the first step in most analyses.

Three challenges remain: correcting strong batch effects, scaling to millions of cells, and modeling how covariates influence gene expression. To address these challenges, we developed PIANO: Probabilistic Inference Autoencoder Networks for multi-Omics, a deep learning framework whose central feature is a generative model of gene expression data. Additionally, PIANO achieves robust integrations and trains 10x faster than previous methods.

PIANO accurately integrates single-cell data across species and across single-cell and spatial transcriptomics modalities. As practical applications, PIANO models spatially-resolved gene expression during Alzheimer’s disease progression in human brains and integrates over 100 million cancer cells to model drug perturbations. In summary, PIANO’s integration and generative modeling capabilities will empower novel insights for countless future studies.

## INTRODUCTION

Single-cell RNA sequencing (scRNAseq) enables the creation of cross-species and cross-modality atlases with millions of cells and nuclei, including comprehensive atlases of mouse and human brains^1,2^. Using this technology, international consortiums, such as the BRAIN Initiative Cell Atlas Network (BICAN) and Human Cell Atlas, are mapping the cellular composition of organs and even entire organisms^3,4^. Comparing data across these studies enables exploring the evolutionary conservation and divergence of cell types across multiple species, developmental timepoints, and modalities^1,2,5,6^. As increasingly large quantities of data are collected, scaling to millions of cells per dataset, there is a growing need for powerful new tools to integrate and analyze these data.

Single-cell data are limited by sparsity and confounding sources of variation — often referred to as batch effects. These batch effects, such as the tissue donor and sequencing platform, distort biological signals and make it difficult to compare cell types across conditions. The goal of performing single-cell integration is to find a more informative, shared representation of the data. This integrated latent representation should remove confounding batch effects while preserving biologically-relevant cell type information.

To perform batch correction, multiple methods have been developed for projecting scRNAseq data to a shared, low-dimensional latent space^7^. One approach is Seurat’s canonical correlation analysis (CCA), which finds pairwise anchors by identifying pairs of similar cells across batches^8^. However, pairwise anchors may over-mix data in situations with imbalanced cell type distributions, and Seurat struggles with scalability to larger datasets. Another popular method, Harmony, uses expectation-maximization (EM) to perform batch correction^7^. However, EM may underperform on imbalanced data with rare or closely related cell type populations. These algorithmic limitations motivate the development of more flexible approaches.

An alternate approach is to find an integrated latent representation using a variational autoencoder (VAE)^9^. A VAE is a non-linear, neural network that utilizes an encoder-decoder architecture to model high-dimensional data using a low-dimensional latent variable. Through the bottleneck imposed by the low-dimensional latent space, the VAE extracts the most meaningful cell type information from the data. The generative model, or decoder, transforms this low-dimensional latent space to the high-dimensional space of gene expression counts. The recognition model, or encoder, enables inference of the latent variable from the observed single cell data. When used for integration, VAEs enable learning a shared latent space across batches. However, most VAE applications in single-cell genomics have focused on the latent space and have largely overlooked the decoder’s capability for generative modeling, which is well-suited for modeling gene expression across experimental conditions, such as disease progression and drug perturbations.

We introduce PIANO: Probabilistic Inference Autoencoder Networks for multi-Omics, a variational autoencoder framework for learning a comprehensive, generative model of single-cell transcriptomics data. PIANO learns robust shared representations across difficult integration regimes, including divergent species, developmental timepoints, deep phylogenetic distances, and data modalities, while preserving cell type identities. Its efficient implementation trains up to 10x faster than previous methods and scales to atlases of 100 million cells. Beyond integration, PIANO’s generative model captures how categorical and continuous covariates shape gene expression counts, enabling numerous applications: correcting counts for batch and species effects, improving the specificity of differential expression, modeling spatial gene expression across Alzheimer’s disease progression, and predicting drug-dose responses in cancer lines. These results establish PIANO as a robust, scalable framework for generative modeling and multi-modal integration of single-cell data, empowering novel insights for future studies.

## RESULTS

### PIANO models single-cell data using a deep generative model

PIANO is a latent variable model for single-cell transcriptomics data. Using a variational autoencoder (VAE)^9^ framework, PIANO uses a recognition model (encoder) to infer the latent variable from gene expression data and a generative model (decoder) to reconstruct the data.

The generative model reconstructs each cell’s gene expression from a low-dimensional latent variable (encoding cell type information) and its covariates. These covariates can be categorical—such as donors, species, sequencing platforms, and modalities—or continuous, such as disease progression or drug dosages. Specifically, the generative model consists of a deep neural network (DNN) and a generalized linear model (GLM). The DNN maps the latent representation 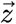 and covariates 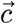 to gene expression proportions *x*_*gene,prop*_. A generalized linear model (GLM) then models the expression of each gene *x*_*μ*_,_*gene*_ under a negative binomial (NB) distribution by performing regression on the gene expression proportion *x*_*gene,prop*_, library size *x*_*L*_(i.e., total gene counts per cell), and covariates 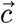:

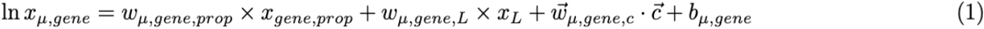

By learning weights for each covariate, PIANO performs generative modeling of gene expression under different scenarios. For example, it can predict changes in gene expression for a given cell under a higher drug dosage (a continuous covariate). For categorical covariates such as donors, which are encoded as one-hot vectors (e.g., 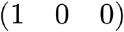), PIANO can utilize a uniform vector (e.g, 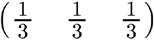) to correct for donor effects. Moreover, by accounting for batch covariates in the generative model, the latent variable is able to prioritize capturing biological variation rather than batch effects.

Exact inference of the latent variable is intractable. Instead, the encoder uses a DNN to infer the latent variable from gene expression data. Due to the low dimensionality of the encoder bottleneck, and since covariates are already accounted for by the generative model, the latent variable is encouraged to learn the most important biological information about each cell rather than learning batch effects. This enables tractable training of the generative model and obtaining integrated latent representations for single-cell data.

Expanding upon the VAE framework, PIANO supports additional features to improve integration performance. PIANO uses a gradient reversal layer^10^ with an adversarial network to further remove batch effects (see methods). Additionally, PIANO can further refine the latent space by using the generative model to obtain batch-corrected counts; using these corrected counts, the encoder can infer a latent space with even stronger batch correction. Together, these design choices enable robust latent space integration.

### PIANO enables scalable performance for single-species integration

PIANO was first benchmarked against publicly available methods on five published single-species datasets (Figure 1B). These datasets varied in the number of cells (16k-373k), cell types (11-81), and tissues, but all contained known batch effects, such as donors and sequencing platforms^11–22^ (Figure 1B). PIANO was benchmarked against widely-used classical methods (Seurat^8,23^, Harmony^7^, and scVI^24^), more recent methods (sysVI^25^ and scDREAMER^26^), and principal component analysis (PCA) — which serves as an unintegrated baseline for these comparisons (Figure 1B). Due to potential stochasticity in model initialization, the deep learning integration methods (PIANO, scVI, sysVI, and scDREAMER) were repeated five times (Table S1). On these datasets, PIANO consistently ranked among the top two methods for overall integration (Figure 1C), with reliable performance across hyperparameters (Table S2).

**Figure 1:**
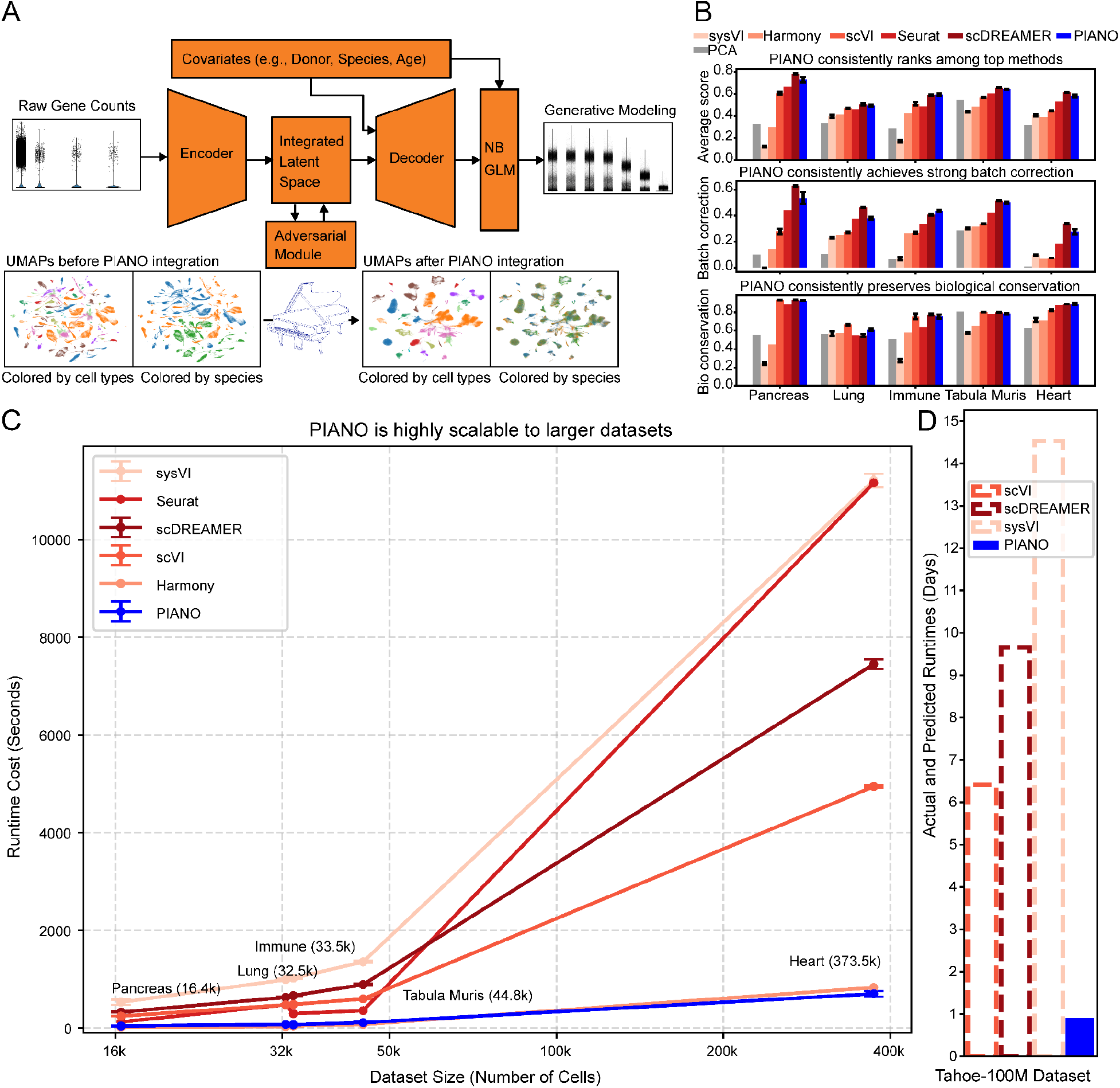
PIANO enables fast, robust, and scalable integration. A) PIANO integrates single-cell RNA transcriptomics data into a probabilistic latent space utilizing a variational inference framework; combined with its negative-binomial (NB) generalized linear model (GLM), PIANO enables generative modeling, such as performing batch correction on gene expression counts, predicting effects of drug perturbations, and modeling disease progression. B) PIANO is consistently a top-performing model on single-species integration tasks. C) PIANO is substantially faster than other deep learning methods. D) PIANO completes training over 100 million cells in just 21.8 hours, whereas other methods are projected to take 1-2 weeks.

Integration performance was quantified using established benchmarking metrics from the scIB package, which quantified the biological conservation (i.e., how well does the latent space preserve known biological cell types) and batch correction (i.e., how well mixed are cells between different batches) of the integration^27^. The average of the normalized mutual information (NMI) and adjusted Rand index (ARI)^27^ was used for quantifying biological conservation. The average of the K-nearest-neighbors batch effect test (KBET) score and integration local inverse Simpson’s index (iLISI) (Methods)^27^ score was used for quantifying batch correction. The overall score was the average between the biological conservation and batch correction scores.

PIANO was also the fastest deep learning method by a wide margin. In comparison to PIANO, scVI took over 7 times longer, scDREAMER took over 10 times longer, and sysVI took almost 16 times longer to train on the larger heart dataset (373k cells). For all deep learning methods, the same hardware was used for training: one NVIDIA A100 GPU with 40Gb of GPU memory. Moreover, for these comparisons, scVI was provided with 11 workers to speed up dataloading — often a training bottleneck — whereas PIANO utilized a single worker for its dataloader. PIANO’s speed and reliability makes it practical for large integrations.

These advantages become especially apparent in truly large atlases. With over 100 million cells, the Tahoe-100M atlas is one of the largest publicly available datasets to date, including not only 50 cancer cell lines but also perturbations over 379 distinct drugs^28^. Previous studies required subsampling to handle the scale of the data^28^. In contrast, using only 21.8 hours of training, PIANO learned a comprehensive generative model of this dataset, utilizing a custom sparse continuous covariate encoding implementation to reduce runtime memory costs (Figure 1D). Assuming the other deep learning methods could withstand the memory demands of this dataset, projections from observed scaling suggest that scVI would take a week to train, scDREAMER over a week and a half, and sysVI would take over two weeks (Figure 1D). These results illustrate PIANO’s suitability to atlas-scale integration.

### PIANO enables cross-species comparisons across primate basal ganglia

Integration across species poses additional challenges because gene expression diverges with increasing evolutionary distance, cell types can be gained or lost in different organisms, and because homologous tissues can still contain dramatically different cell type proportions. Ideally, integration should correct batch effects (such as within-species donor effects) while still appropriately mixing cognate cell types across species and preserving biological cell types. To better understand the capabilities of VAE models for cross-species integrations, multiple approaches were investigated for feature selection and model architecture using a large, multi-species brain cell atlas of the primate basal ganglia^29,30^. The Human and Mammalian Brain Atlas (HMBA) – part of BICAN – contains RNA sequencing data from nearly 2 million single nuclei sampled across human, macaque, and marmoset basal ganglia structures^29,30^. Each species’ data were originally computationally integrated using scVI to generate a consensus cross-species cell taxonomy at two hierarchical levels: cell Class (12 Classes) and cell Group (61 Groups)^29^. These data provide a valuable resource for cross-species comparisons.

To balance the number of cells between species, up to 6,400 cells per Group were included for each species. Using the Group labels to assess biological conservation, PIANO achieved both better separation of cell types (biological conservation) and mixing between donors and species (batch correction) compared to scVI and scDREAMER (Figure 2A, Table S3). PIANO also trained substantially faster than both scVI and scDREAMER (Figure 2B). These comparisons used the same architecture, with 3 hidden layers, 256 hidden nodes, and 32 latent dimensions for each method, which has a well-balanced performance (Table S4).

**Figure 2:**
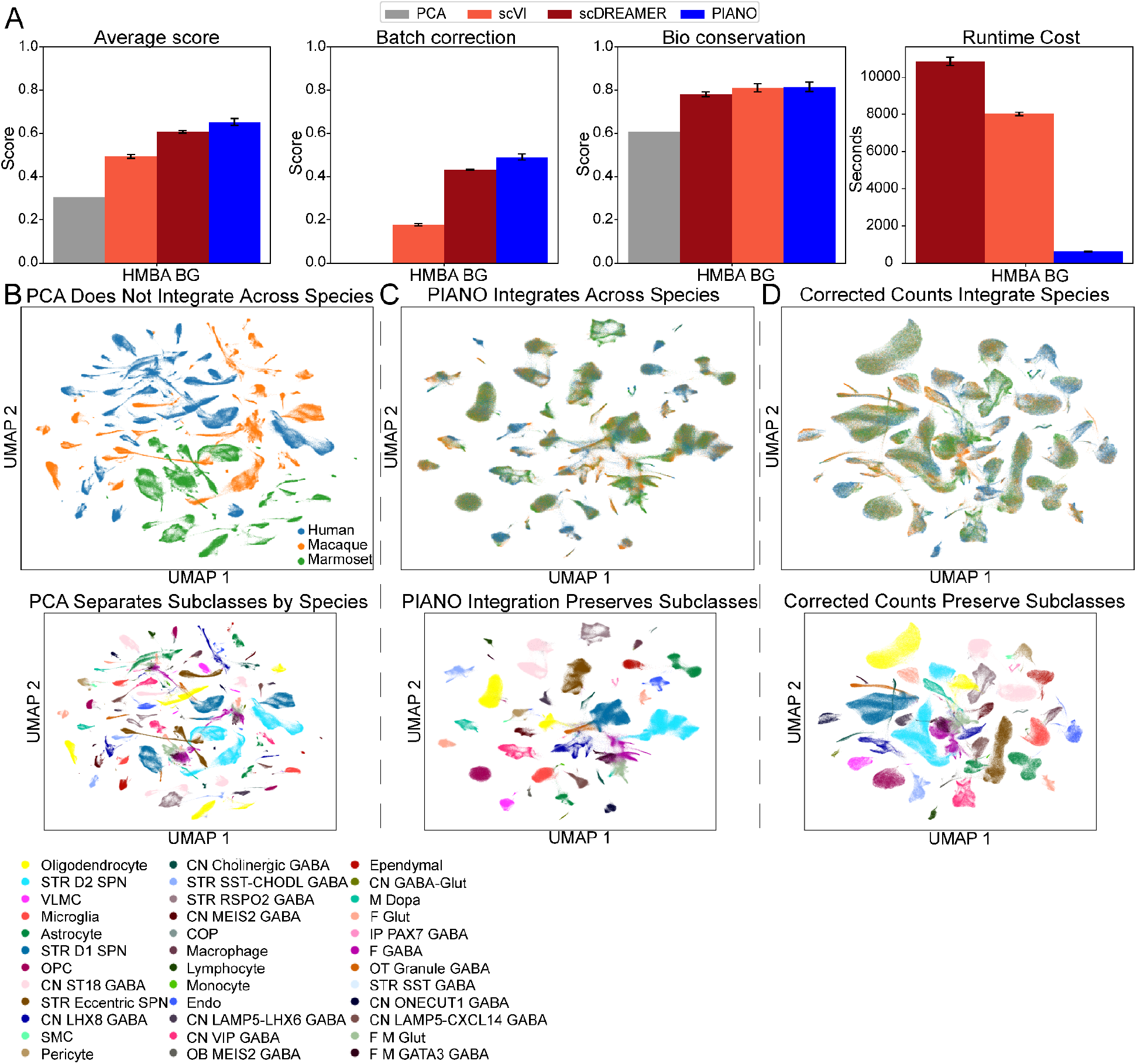
Cross-species integration of primate basal ganglia. A) PIANO enables superior biological conservation of cell types and stronger batch correction compared to the top performing deep learning methods while requiring a fraction of the runtime costs. B) Principal component analysis (PCA) was performed on the primate basal ganglia gene expression counts and projected into two dimensions using Uniform Manifold Approximation and Projection (UMAP). The UMAPs are colored by species (top) and Subclass (bottom). Cell types that should be conserved across species (mixed) are instead shown in separate clusters, indicating a lack of integration across species using PCA. C) PIANO’s integration and generative modeling is used to correct counts across donors and species and then retrieve a latent representation. TheUMAP shows robust integration, where conserved cell types are mixed across species (top) and distinct cell types remain separated (bottom). D) PIANO enables gene expression counts correction for batch effects. Unlike the PCA representation of the original counts data in A), the PCA representation of the PIANO-corrected counts shows integration across species (top) while preserving cell type clusters (bottom).

Although PCA had reasonable biological conservation, it offered no batch correction between species, demonstrating the necessity of integration methods for cross-species comparisons (Figure 2C). PIANO not only obtained a robust integrated latent representation (Figure 2D), but its generative model also enabled correcting gene expression counts for donor and species effects. In fact, using these corrected counts, PCA was able to find an integrated latent space via linear dimensionality reduction—with strong batch (species) correction and preservation of cell types—demonstrating the robustness of the counts correction (Figure 2E). These results showed that PIANO both corrects gene expression counts for batch effects and produces a robust latent space integration.

#### PIANO’s integration is robust to gene selection

Feature selection is an important consideration for cross-species integration. Prior work showed that functional gene sets, such as transcription factors, can discriminate between cell types nearly as well as all expressed genes^2^. Using the HMBA data, 4,096 (2^12^) highly variable genes (HVG) using the Seurat v3^8,31^ approach with species as the batch key were compared against functional gene sets (transcription factors, synapse-related genes, and extracellular-matrix-related genes) as well as 4,096 randomly selected genes (Figure S1). Larger gene sets performed comparably regardless of functional category, and even random genes generally matched HVGs apart from slightly lower biological conservation. PIANO’s integration is therefore robust to the choice of feature set and is more sensitive to the number of genes than on their annotation.

#### *In silico* perturbation as a test of species-specific cell type preservation

One mode of evolutionary divergence could come as complete gain or loss of a cell type^32^. Ideally, cross-species integrations should preserve species-specific cell types^33^. As a robustness validation, two *in silico* perturbation experiments were performed on evolutionarily conserved cell types. To determine if a species-specific cell type would be merged into a related cell type shared across species, only the eccentric Spiny Projection Neuron (eSPN) Subclass was first withheld from the non-human primate datasets, such that only the human basal ganglia dataset retained this Subclass. In this scenario, both PIANO and scVI kept the human-specific eSPNs separate from the related D1 and D2 SPNs, which were still shared across primates (Figure S2A-B). This demonstrates that VAE integration can preserve novel cell types.

Next, only the non-human primate D1 SPNs were withheld. Since the D1 SPNs are far more transcriptionally similar to the D2 SPNs than are the eSPNs, this constituted a more difficult perturbation. Although both scVI and PIANO initially mixed the D1 and D2 SPNs, by turning off adversarial training and using the original latent space (without correcting raw counts for global species effects), PIANO successfully separated the human-specific D1 SPNs from the shared D2 SPNs (Figure S2C-D). This showed that gene counts correction should be carefully applied to avoid global overcorrection, such as by checking that the latent space does not merge cell types that were separated in the single-species latent space. In such scenarios, count correction should be reserved for cell types that are conserved across species.

#### Differential gene expression analysis between human and non-human primates

A common approach to examining how specific cell types diverge across species is to analyze their differentially expressed genes (DEGs). However, gene expression counts are noisy and influenced by donor and global species effects. To mitigate these effects, PIANO explicitly models donor and species effects, which improves the robustness of DEG analyses.

Cross-species DEG comparisons for each Subclass were first performed between humans and non-human primates—correcting for both donor and species effects—to examine cell type specific divergences (controlling for global species effects). Here, PIANO’s corrected counts approach was more stringent on the number of calculated DEGs compared to the number of DEGs calculated using raw counts, reducing false positives (Figure S3). In particular, a substantial proportion of the DEGs excluded were global species DEGs, which enhances the cell type specificity of DEGs (Figure S3). The non-neuronal cell types in these cross-species comparisons had the most numerous DEGs, suggesting stronger evolutionary divergence in non-neuronal brain cell types than in neuronal brain cell types^34^ (Figure S3).

The previous analysis found DEGs that were species-specific; in other contexts, it is useful to know which genes are species-conserved DEGs amongst species. The DEGs that uniquely and commonly define a conserved cell type were examined. The DEGs between each pair of Subclasses were computed, the genes that had higher expression in a homologous Subclass over all pairs of cell type comparisons for all three species (human, macaque, and marmoset) were selected. In these comparisons, PIANO reduced the effects of outliers in single-cell data, recovering DEGs that would be misconstrued as human-specific but are actually conserved across primates (Figure S4A). For example, in marmosets, the *CRYAB* gene was not detected as a DEG in oligodendrocytes due to a small number of outliers with high expression in non-oligodendrocytes (Figure S4B). These analyses showed that modeling donor and species effects may improve specificity for cross-species DEG analysis, reducing false-positive and outlier-driven errors when assessing cell type conservation and divergence.

### PIANO enables integration across species and developmental timepoints

Cell type identities are especially dynamic during maturation^33^. This challenge is amplified when attempting to integrate across species, since developmental timepoints may not be aligned. In a recent study, astrocyte development across embryonic and postnatal timepoints in mice and marmosets was compared but was not successfully integrated across species^33^.

PIANO obtained robust integrations of the complete dataset across timepoints, including both neurons and non-neuronal cells, outperforming both scVI and scDREAMER for integration performance and runtime (Figure 3A). PIANO obtained strong integration with good mixing across species (Figure 3B). Additionally, distinct region differences observed in both neurons and glia were preserved (Figure 3C), recapitulating region specificity of glial cell types as described in the original manuscript^33^. Moreover, PIANO separated certain developmental time points—such as cells in the embryonic stage distinctly separated from the clusters in the later time points (Figure 3D)—which were mixed together in the previous single species integrations^33^. The original manuscript also identified two species-specific neuronal superclusters: an embryonic mouse type ‘Inh Str Immature’, and a marmoset thalamic GABAergic type ‘Inh Thal MB-der’; PIANO preserved both species-specific cell types (Figure 3E). These findings showed that in a single analysis, PIANO integrated evolutionary-distant species while preserving signals from regional and developmental sources.

**Figure 3:**
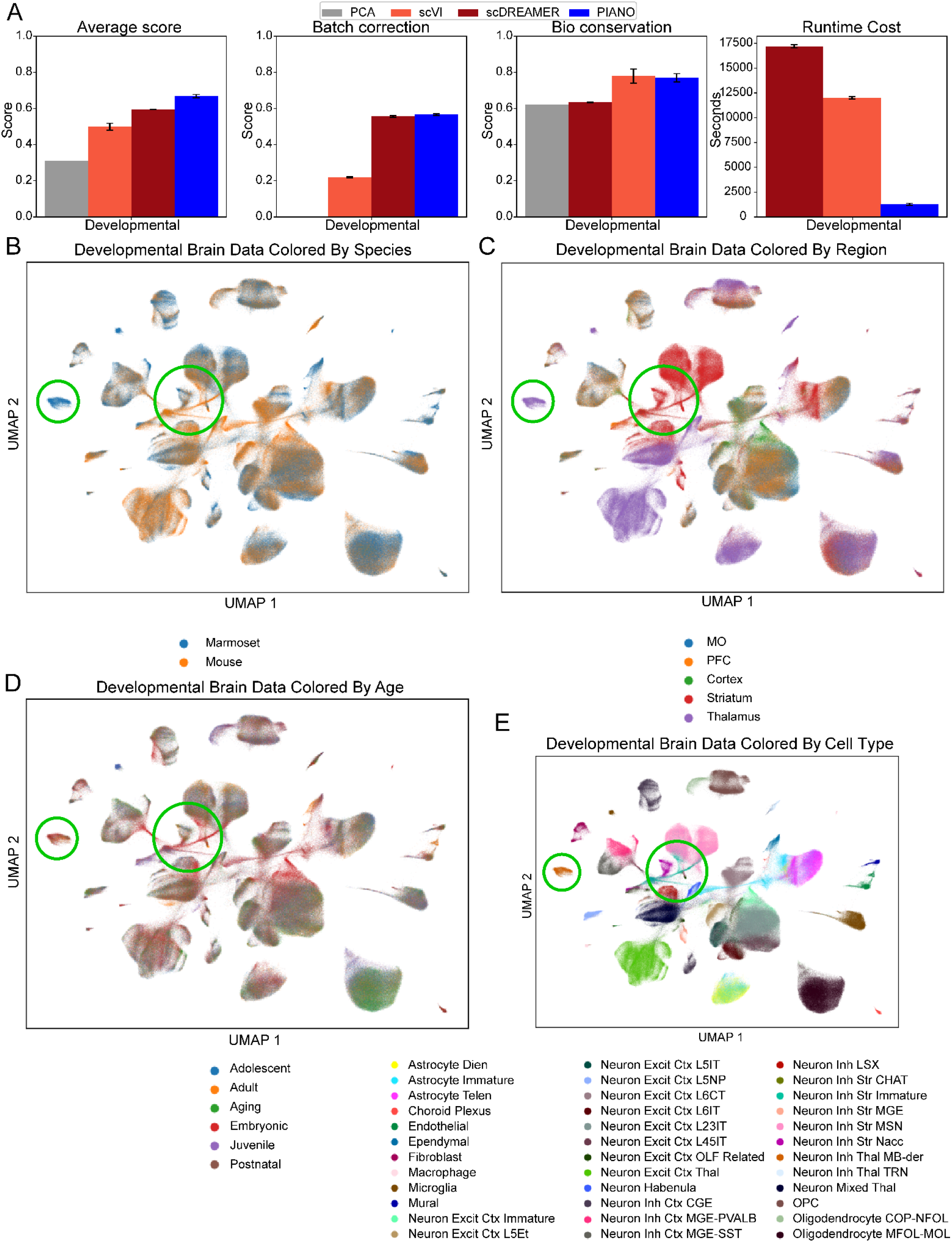
Cross-species developmental brain data. A) PIANO achieves robust integration performance with substantially faster runtime compared to scVI and scDREAMER. PIANO has substantially stronger batch correction than scVI with similar biological conservation of cell types and substantially stronger biological conservation than scDREAMER with similar batch correction. B-C) UMAP representation of PIANO integration shows robust cross-species mixing across multiple brain regions. D-E) PIANO integration preserves cell type identities, including a marmoset-specific inhibitory thalamic type (smaller circle) and a developmental, mouse-specific inhibitory neuronal type (larger circle).

In comparison, scVI had similar separation of cell types but substantially less mixing across species (Figure 3A). Although scDREAMER had similar batch correction to PIANO and stronger batch correction than scVI, it had lower biological conservation and substantially longer runtime costs (Figure 3A). Furthermore, scDREAMER suffered from training instability: only two out of five runs were able to train without crashing due to numerical instability (Table S5).

### PIANO resolves shared and species-specific dynamics of differentiation trajectories across deep evolutionary distances

Cross-species comparative analyses of single-cell atlases provide a powerful framework for decoding evolutionary dynamics of cell-types and characterizing cellular diversity in understudied lineages. However, as taxon sampling in single-cell atlases becomes wider, constructing common references is increasingly challenging because orthology relationships among genes become more complex. To evaluate PIANO’s performance in this setting, we applied it to a testis single-cell dataset spanning 11 vertebrate species across broad phylogenetic distances^35^. Previous work reported that other methods relying primarily on one-to-one orthologs perform poorly in this dataset, whereas approaches that incorporate many-to-many homologous genes are computationally demanding and, in some cases, fail to run because of memory requirements^36^.

PIANO successfully integrated cells across the 7 primates as well as the rodent, marsupial, monotreme, and avian lineages represented in the dataset (Figure 4A). Importantly, the integrated space retained the expected structure of spermatogenesis: germ cells formed a continuous differentiation trajectory, while the annotated major cell types remained distinguishable (Figure 4B). This indicates that PIANO preserves both shared developmental structure and cell-type identity.

**Figure 4:**
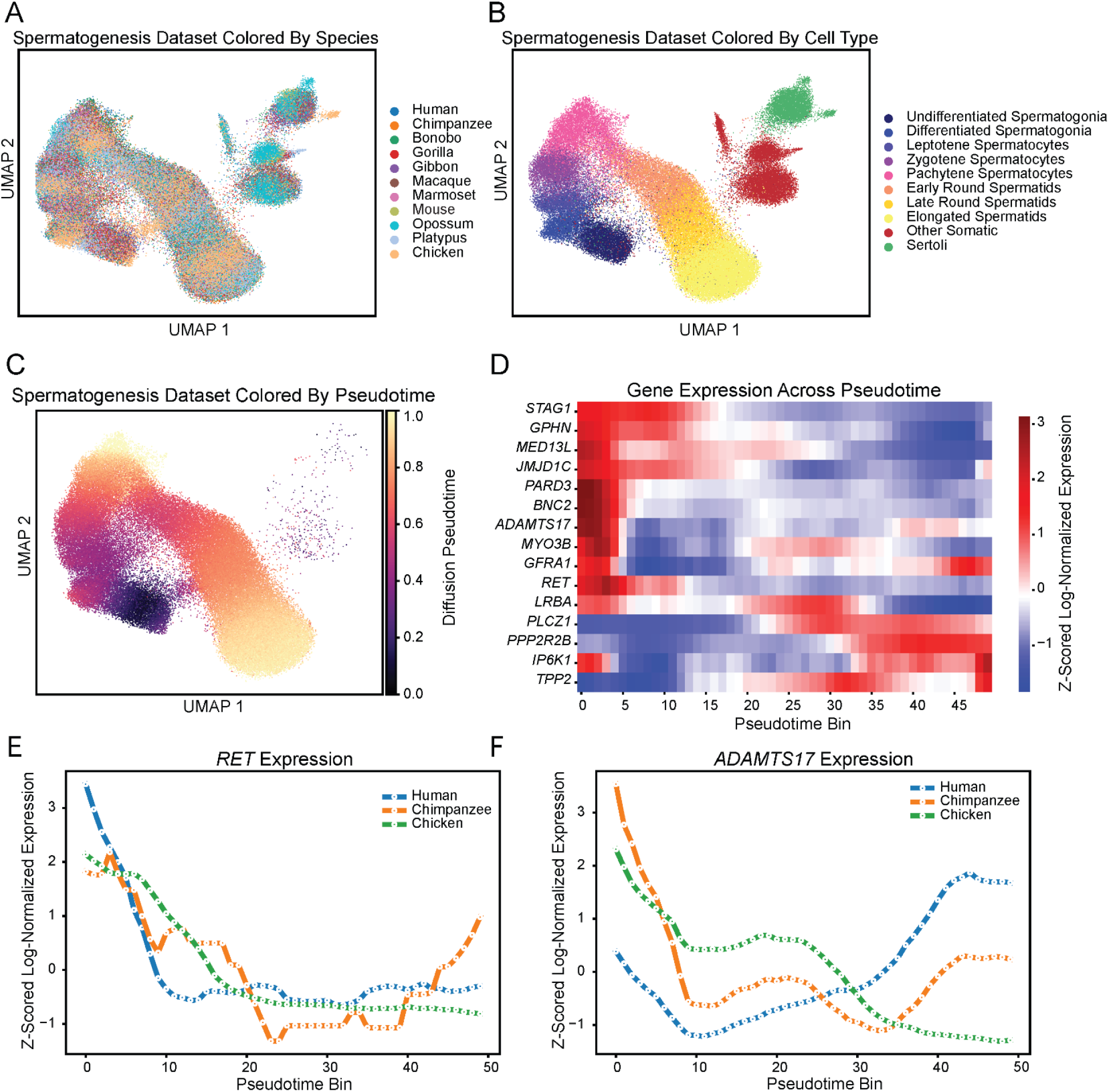
PIANO captures a conserved axis of spermatogenic differentiation across vertebrates. A) PIANO achieves robust cross-species integration across vertebrates B) while preserving testis cell-type identities. C) Pseudotime of spermatogenic cell progression overlaid on PIANO UMAP projection. Pseudotime values were capped at the 99th percentile to reduce the influence of outliers and then rescaled from 0 to 1 for visualization. D) Heatmap of Z-scored log-normalized expression for representative genes with dynamic expression profiles across pseudotime bins. E-F) Mean expression profiles across pseudotimes bins for Human, Chimpanzee and Chicken. *RET* and *ADAMTS17* are illustrative examples of conserved and shifted trajectories, respectively.

To further assess whether the PIANO latent space captured meaningful continuous developmental structure, pseudotime was inferred among germ cells after excluding somatic cell populations, including Sertoli cells. Diffusion pseudotime was computed from a neighborhood graph constructed using the integrated PIANO latent representation. A clear progression was observed from undifferentiated spermatogonia to the pachytene spermatocytes and then from early round spermatids to the final stage of elongated spermatids (Figure 4C). Among these trajectories, multiple genes displayed known transcribed markers of germ cell differentiation, such as *GFRA1*, a known receptor marker for undifferentiated spermatogonia^37^ and *TPP2*, a tubulin polymerization promoting gene which is involved in the formation and morphology of spermatids^38^ (Figure 4D). The shared pseudotime axis was then used to compare gene expression dynamics across species. This analysis recovered conserved trajectories, such as *RET*, as well as genes with species-specific shifts, such as *ADAMTS17*, a protease whose increased expression in human spermatids has previously been validated by *in situ* hybridization^35^ (Figure 4E-F). Together, the resulting PIANO latent space recovered a shared spermatogenesis trajectory, enabling systematic comparisons of conserved developmental progression, species-specific deviations and the evolution of gene programs across deeply diverged vertebrate lineages. Thus, a single PIANO model resolved a conserved spermatogenic trajectory across deep evolutionary distances while distinguishing conserved gene programs (e.g., *RET*) from lineage-specific shifts (e.g., *ADAMTS17*).

### PIANO enables multi-modal data integration with spatial transcriptomics

Along with integrating datasets across multiple species and timepoints, integration across data modalities further empowers biological discovery. Spatial transcriptomics technologies, such as Multiplexed Error-Robust Fluorescence *In Situ* Hybridization (MERFISH)^39^, enable identifying the exact physical locations of cells in a slice of tissue, which is lost in single-cell data; however, this spatial resolution comes at the cost of capturing far fewer genes and/or transcripts. Recently, the Broad Institute utilized MERFISH to create a whole mouse brain spatial atlas containing a complete spatial transcriptomics series of four mouse brains^40^; as it contained more genes, the Broad Institute’s MERFISH data with 1,122 genes shared across gene panels^40^ was used instead of the Allen Institute’s MERFISH data with a 500 gene panel^2^ to demonstrate single-cell and spatial transcriptomics integration with PIANO.

To obtain cell type labels, the Broad Institute spatial atlas was originally integrated and mapped to the Allen Institute’s whole mouse brain single cell atlas^2^ using Seurat (CCA)^40,41^. Given the stronger performance of PIANO and scVI relative to Seurat, the integration of the Broad Institute’s whole mouse brain spatial transcriptomics atlas with the Allen Institute’s whole mouse brain single cell transcriptomic atlas was revisited using these two methods, excluding scDREAMER due to its longer runtime costs, training instability on complex datasets, and tendency to lose cell type resolution. The genes in the single cell transcriptomics data and spatial transcriptomics data were first subset to the 1,122 genes shared in all datasets^2,40^; the cells in each datasets were then subset to up to 6400 cells per Subclass to obtain a balanced dataset for integration. Comparing PIANO to scVI, similar biological conservation (separation of ~300 Subclass resolution cell types) and substantially stronger batch correction (across data modalities) was observed (Figure 5A, Table S6). Despite the complexity of the dataset, PIANO obtained robust separation of the cell types (Figure 5B) and good mixing across data modalities (Figure 5C) and sequencing platforms (Figure 5D). These results demonstrated that PIANO reliably enables cross-modality integration between single-cell and spatial transcriptomics data.

**Figure 5:**
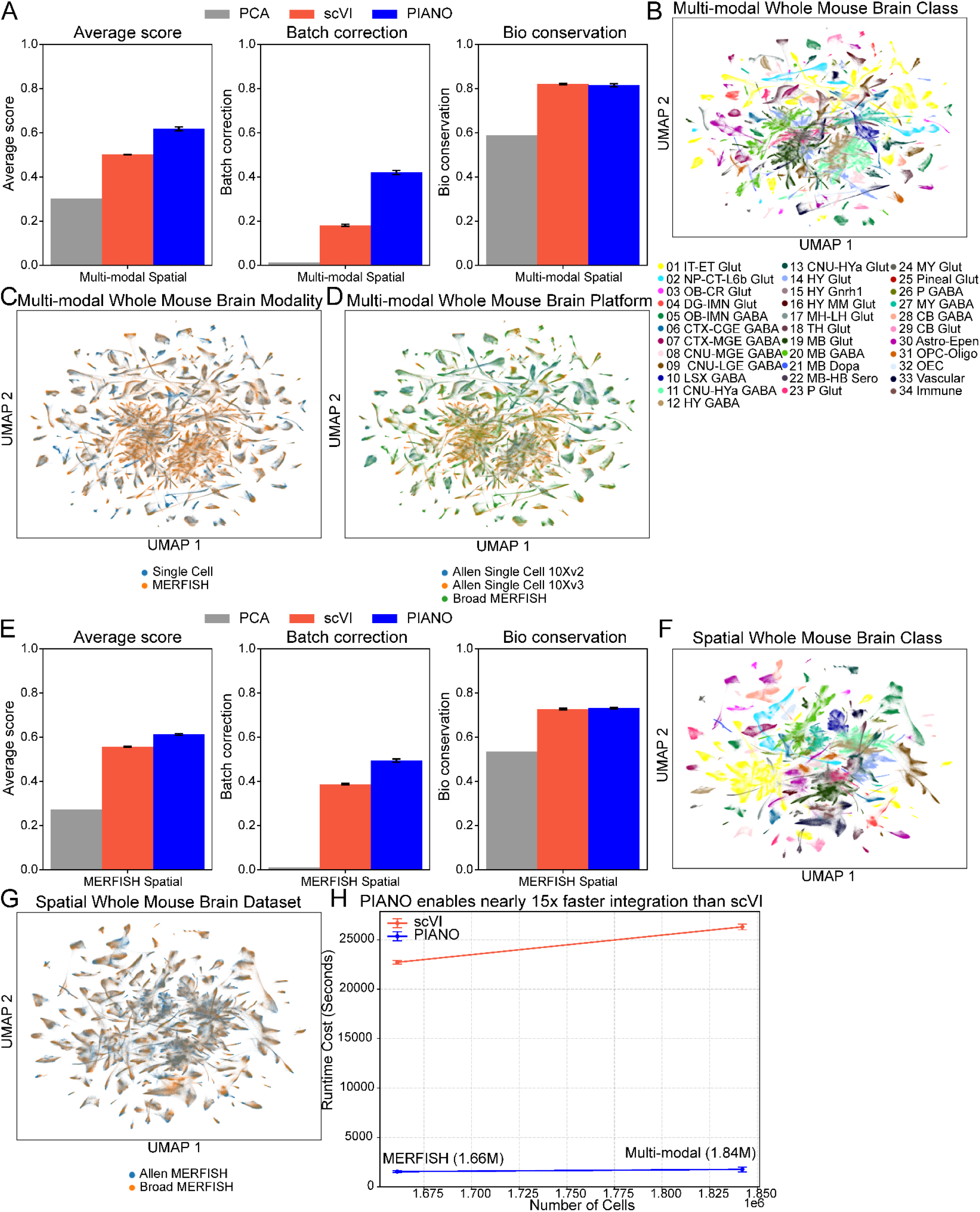
Multi-modal integration of whole mouse brain. A) PIANO enables superior integration of whole mouse brain data between single-cell and spatial transcriptomics (MERFISH) modalities than scVI, with similar biological conservation of cell types. B-D) Within each Class-level cell type, PIANO separates the Subclasses (not colored due to the large number of labels) while integrating across modalities and platforms. E-G) PIANO achieves similar biological conservation of Subclasses and far stronger batch correction for integration across multiple MERFISH datasets, preserving cell types while integrating across datasets. H) PIANO achieves these superior integrations with nearly 15x faster training compared to scVI.

Additionally, PIANO integration was investigated across multiple spatial transcriptomics datasets. Combining spatial datasets can improve generalizability across donors and expand spatial coverage, such as with different sampling across brain regions or by combining coronal and sagittal brain slices. Using two separate MERFISH datasets—one from the Broad Institute and the other from Allen Institute whole mouse brain atlases—the data was subset to the shared 477 genes across gene panels. Despite having an even smaller set of genes available for integration than the single-cell and spatial integration, PIANO still obtained robust integration across datasets with similar biological conservation of Subclass resolution cell types to scVI and superior mixing across datasets (Figure 5E, Table S6). Again, despite having fewer genes, PIANO still maintained separation of cell types (Figure 5F) while mixing appropriately across datasets (Figure 5G) Moreover, for both the spatial transcriptomics and multi-modal integrations, PIANO trained substantially faster than scVI, requiring less than 7% of the runtime costs (Figure 5H, Table S6), demonstrating PIANO’s scalability to larger datasets.

### PIANO enables spatial modeling of Alzheimer’s disease progression

Studies using postmortem samples from human donors are extremely valuable, but any postmortem sample is necessarily restricted to a single timepoint. PIANO’s multi-modal integration and generative modeling capabilities were applied to examine the progression of neurodegenerative disease phenotypes. The Seattle Alzheimer Disease Brain Cell Atlas (SEA-AD) middle temporal gyrus (MTG) dataset was selected for this example, which included over 1.1 million nuclei from 84 donors and over 1.5 million spatial transcriptomic cells (spanning 69 sections from 27 donors) profiled using Multiplexed Error-Robust Fluorescence *In Situ* Hybridization (MERFISH). The study authors constructed a hierarchical cell type taxonomy consisting of Class, Subclass, and Supertype^42^. Using neuropathological data common to AD progression, the authors constructed a continuous pseudo-progression score (CPS) for each donor (0 to 1)^42^, which was used as a proxy for disease progression in our analysis. A generative model was learned across these cell types, using CPS to explicitly model Alzheimer’s disease progression trends under that proxy. Donor ID and sex are included as categorical covariates, while age at death and CPS are modeled as continuous covariates. Trained on the scRNA-seq data, PIANO produced a robust representation across subclasses (Figure 6A) and used CPS to model gene expression across all disease progression (Methods).

**Figure 6:**
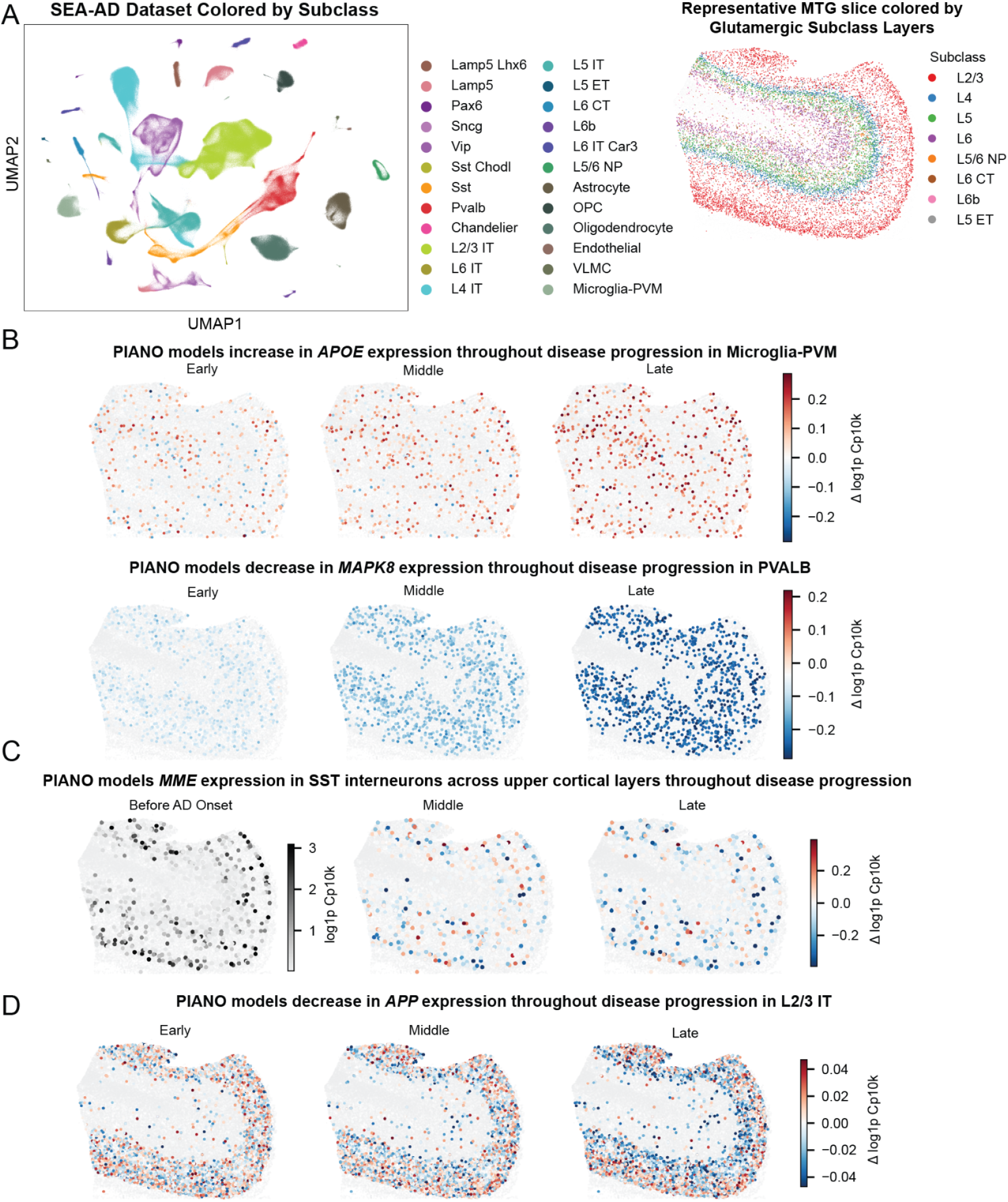
PIANO enables spatial projection of in-silico generated modeled gene expression. A) PIANO achieves robust integration across subclasses using generative modeling, shown by UMAP of the SEA-AD dataset colored by subclass. Glutamatergic cortical layer organization projected onto a MERFISH slice. B) Spatial projection of generated gene expression onto a MERFISH slice across Early (0.2 CPS), Middle (0.5 CPS), and Late (1.0 CPS) disease stages, shown as a difference relative to baseline (0.0 CPS), revealing an increase in *APOE* and decreases in *MAPK8*. D-E) Spatial projection of *MME* and APP expression in upper cortical layers across Before Onset, Early Middle, and Late disease stages, revealing a layer-specific MME decrease in SST cells and shifting *APP* expression between upper cortical layers (increase in L2, decrease in L3).

PIANO used the spatial and snRNA-seq datasets to model spatial gene expression during disease progression. The original authors plotted gene expression across CPS with the single-nucleus data, capturing changes in cellular composition across disease and confirming the spatial distribution of cell type populations. A recent preprint extended the SEA-AD atlas to ten neocortical and allocortical regions, profiling over seven million cells with single-nucleus and spatial transcriptomics to map transcriptional and compositional changes^43^. However, neither study fully bridged the two modalities to model spatially-resolved, transcriptome-wide gene expression changes during disease progression.

The MTG MERFISH data resolved cells in their spatial context. However, the panel contained only 140 genes, which misses numerous disease-relevant genes. To meet this challenge, PIANO was used to integrate the single-nucleus data with the spatial transcriptomics data, mapping the molecular profiles of nuclei to spatial coordinates using K-nearest neighbors (see Methods). Spatially-resolved transcriptomic changes during disease progression were visualized on a representative MERFISH cortical slice (Figure 6B), enabling disease-associated gene expression viewed within a consistent spatial context. Illustrative examples (*APOE* and *MAPK8*) demonstrated transcriptional changes relative to before the onset of disease.

Importantly, this multi-modal integrative analysis revealed layer-specific gene expression patterns that would otherwise be difficult to observe with either modality alone. The authors of the original study conducted cell type composition analysis by comparing the relative abundance (cell proportions) of Supertypes along the CPS, with cell types that decrease in their relative abundance classified as “vulnerable”. The authors observed that vulnerable SST Supertypes exhibited a decrease in *MME* expression^42^. More broadly, PIANO’s generative modeling revealed that a decrease in *MME* was observed across all upper cortical layer SST interneurons (Figure 6D). Expression of *MME* (and decrease during disease progression) was observed in the upper cortical layers, regardless of the status as a vulnerable or unaffected (Figure S5B).

Moreover, a layer-specific pattern was uncovered in amyloid-β related gene expression, such as the Amyloid precursor protein (*APP*). The *APP* expression in L2/3 IT projection neurons across disease progression separated into two subsets. The L2/3 IT neurons that populated L2 showed increased *APP* expression across CPS, while L2/3 IT neurons that populated L3 showed decreased *APP* expression across CPS (Figure 6D). These changes were largely driven by L2 specific Supertypes (L2/3 IT_1, L2/3 IT_6) and L3 Supertypes (L2/3 IT_10, L2/3 IT_13) (Figure S5C), where *APP* expression in the Supertypes specific to L2 are selectively increasing across disease progression, while L3 Supertypes are decreasing (Figure S5D). Together, these results demonstrate PIANO’s ability to model gene expression across disease progression, project these dynamics onto their spatial context, and reveal spatiotemporal patterns that are challenging to resolve using either modality in isolation.

### PIANO enables modeling drug perturbations over 100 million cancer cells

Single-cell transcriptomics data can measure changes in gene expression induced by drug perturbations. The Tahoe-100M dataset provides a publicly-available resource containing 100 million cells from 50 cancer cell lines with 379 unique drugs^28^.

Beyond its size, these data pose additional challenges for modeling drug perturbations. The data are highly sparse, with only 5.3% non-zero entries, and after subsetting the data to the top 4,096 highly variable genes, the unique transcripts per cell contained a mean of only 440.4 counts and a median of only 331 counts. Although the dataset contains a large total number of cells, these cells are distributed across 50 cell lines and hundreds of unique drugs, yielding modest sample sizes for any specific cell line and drug combination. Moreover, each drug was administered with only three non-control dosages, which limits identification of dose-response relationships. These technical challenges reduce the signal available in the Tahoe-100M data.

PIANO presents a practical solution to these challenges. With its speed and scalability, PIANO learns a comprehensive model across the entire dataset, without being forced to subset the data due to runtime or memory limitations (Figure 1D, Methods). This enables PIANO to benefit from pattern recognition across all of the cells and drug perturbations (Figure 7). Using these patterns, PIANO enables generative modeling of how gene expression in each cell line is influenced by drug perturbations.

**Figure 7:**
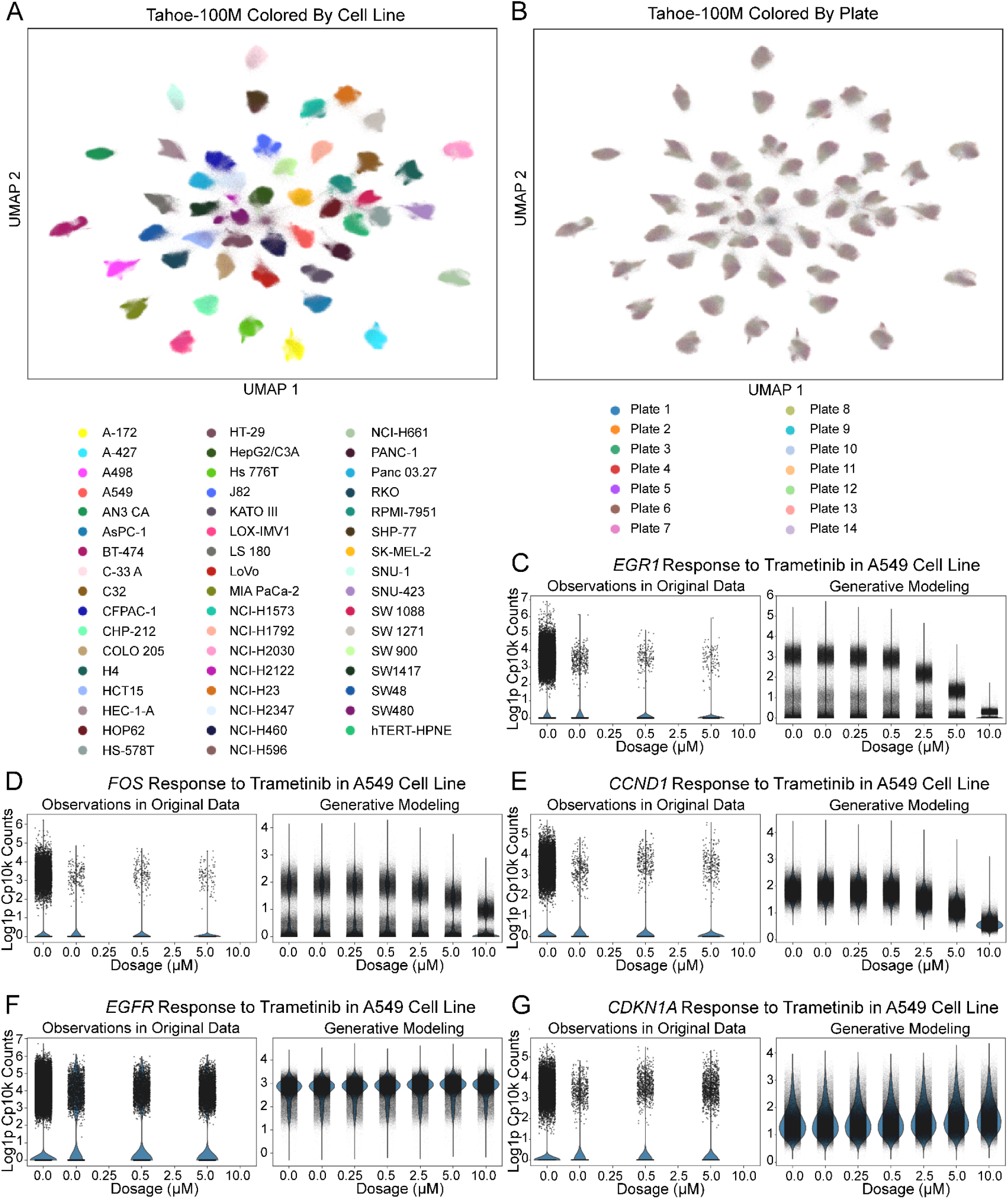
Atlas-scale integration of over 100 million cancer cells. A-B) PIANO enables robust separation of cell lines and mixing across the 14 plates on which the cells were grown. C-E) PIANO recapitulates the expected decrease in transcription of immediate early genes *EGR1* and *FOS* and cell cycle gene *CCND1* in the A549 lung cancer cell line due to treatment with the *MEK* inhibitor Trametinib. F-G) PIANO predicts stable expression of the *EGFR* gene, which is upstream of *MEK*, and stable expression of *CDKN1A*, which competes with *CCND1* to prevent cell proliferation^46^ (relevant to cancer progression).

As an illustrative example, changes in gene expression resulting from increasing dosages of Trametinib were examined in the commonly studied A549 lung cancer cell line^44^. The A549 cell line features an oncogenic mutation in the *KRAS* gene^44^. The resulting kinases initiate a well-known cell signaling cascade through the *RAF, MEK*, and *ERK* kinases, which make this pathway a desirable target for therapeutic treatments^44,45^. One such treatment is Trametinib, which is a robust *MEK* inhibitor^44^. Trametinib inhibits *MEK* from activating *ERK*, which would have otherwise activated the downstream *ERK* pathway^46^. This pathway upregulates the transcription of immediate early genes, such as *FOS* and *EGR1*, and key cell-cycle genes, such as *CCND1*, which enables cell proliferation^46^.

Although these pathways are well understood, the Tahoe-100M dataset is sparse and only provides 3 dosages per drug perturbation, making it difficult to identify trends in the original gene expression data (Figure 7C-G). PIANO recovered the signal with its generative modeling of gene expression counts across simulated drug perturbation dosages (Figure 7C-G). As expected, PIANO recovered decreases in gene expression of the *FOS, EGR1*, and *CCND1* genes with increasing Trametinib dosage (Figure 7C-E). Moreover, genes that are not expected to change in expression due to this perturbation, such as *EGFR* (an upstream, receptor tyrosine kinase)^46^ maintained their stable, consistent gene expression levels (Figure 7F) across the simulated dosages. Similarly, *CDKN1A*, a cell-cycle gene that competes with *CCND1* to ensure the cell does not proliferate^46^, maintained consistent levels of gene expression (Figure 7F). These results demonstrate PIANO’s capability to recover biological signals from noisy data.

## DISCUSSION

### PIANO enables fast reliable integrations and robust generative modeling

PIANO is a framework for generative modeling of single-cell multi-omics data. By combining a latent variable model with a regression model, PIANO uses the latent variable to extract cell type information while explaining donor and species effects using covariates, thereby performing batch correction. Additionally, PIANO learns how continuous covariates influence gene expression, which enables modeling drug perturbations and neurodegenerative disease. Moreover, PIANO is fast and scalable to atlas-sized datasets with 100 million cells. We perform PIANO integrations on datasets encompassing multiple species, developmental timepoints, data modalities, and disease conditions. These demonstrate PIANO’s utility for common analyses – latent space integration, differential gene expression, and pseudotime trajectories – that characterize conserved and diverged biology across cell types.

The comparative integrations shown in the present work illustrate how PIANO can reveal biological structure that is otherwise difficult to resolve across species. In the primate basal ganglia, modeling donor- and species-effects recovered conserved cell-type markers that raw counts would have misclassified as human-specific. In the developing brain, PIANO aligned mouse and marmoset datasets that previous methods could not, while preserving species- and stage-specific populations. Moreover, across eleven vertebrates spanning deep evolutionary distance, a single PIANO model resolved a conserved spermatogenic trajectory, distinguishing conserved gene programs (e.g., *RET*) from lineage-specific shifts (e.g., *ADAMTS17*).

Along with robust integrations, PIANO is designed to meet the demands of atlas-scale datasets. Compared with other implementations that may utilize multiple GPUs—which are both multiplicatively more expensive to use (scaling with the number of GPUs used) and introduce additional runtime overhead costs (e.g., for inter-device communication and code that cannot be parallelized)—PIANO’s light-weight implementation is faster and more cost-efficient. Moreover, modern atlases are often so large that they are stored in separate AnnData objects^47^, and concatenating multiple AnnDatas together for training is computationally expensive. PIANO natively supports training on multiple AnnData objects, which makes PIANO easy to use for new integrations—especially for ongoing data collection.

PIANO explicitly models how covariates influence gene expression, which enables generative modeling of gene expression under different scenarios, such as neurodegenerative disease. Along with recapitulating previous findings on Alzheimer’s disease, PIANO enables novel spatial modeling of gene expression patterns. Studying disease progression is challenging due to post-mortem samples being limited to their corresponding ages, time points, and disease conditions, along with donor to donor variability. PIANO’s integrative analysis across donors and disease time points enables learning a robust, comprehensive generative model of disease progression. This integrative approach unlocks new capabilities, such as modeling genes found only in the single-nucleus data and not the spatial panel, including the layer-specific expression of *MME* in SST neurons and the laminar boundary between increases and decreases in expression of *APP* during disease progression. These examples demonstrate how PIANO enables novel multifaceted analyses.

Furthermore, PIANO models truly large atlases with over 100 million cells, recapitulating established cancer biology^44–46^ and unlocking novel analyses of the Tahoe-100M dataset. With higher-resolution analyses compared to the cell-cluster distance and gene set approaches used in the original Tahoe-100M paper^28^, PIANO enables robust generative modeling of how individual genes respond to drug perturbations, revealing patterns that were obscured by the sparsity of the raw data. These results highlight how PIANO’s generative modeling capabilities can empower future studies.

### Data quantity and quality influences integration results

Dramatic differences in dataset quantity and quality can affect the quality of integration results. Although PIANO had reliable performance on the single-species datasets, the advantages of using PIANO integration became more apparent on the larger and higher quality brain datasets, demonstrating substantial performance improvements with both integration and training speed over previous methods. This reinforces the value of high-quality sequencing data to distinguish between evolutionary divergences across species with confounding noise artifacts. The largest dataset considered, the Tahoe-100M dataset, had a low quality-control threshold of only requiring 700 unique molecular identifiers (gene counts) per cell in the raw data^28^, which was further reduced after feature selection subsetting to the top 4096 highly variable genes. Despite having extremely high sparsity, low counts per cell, and limited samples per drug perturbation, the Tahoe-100M dataset had such a vast quantity of total cells that it allowed PIANO to learn how to recognize patterns across the entirety of the dataset, retrieving rich biological signals from substantial technical noise.

Just as the Seurat method was inspired by pointillism painting^23^, the name of the our method, PIANO, serves as a metaphor between integrating complex atlases to performing piano concertos, such as Rachmaninoff’s Piano Concerto No. 2 in C minor^23,48^. Analogous to how the quality of the performance depends on individual contributions of the pianist, conductor, and orchestra, the success of complex integrations relies on having large quantities of high-quality data. A successful performance in lieu of prodigious musical talent requires countless hours of dedicated practice, similar to how the contributions from over 100 million individual cells in the Tahoe-100M dataset overcame its sparsity and low sequencing depth.

### Gene selection

Previous work emphasized functional gene sets for preserving cell type identities^2^. The Allen Institute whole mouse brain (WMB) atlas compared cross-validation accuracy of clustering using all cluster-wise differentially expressed genes (n=8,460 genes) against several functional gene sets including transcription factors (TFs, n=534 genes), cell adhesion molecules (n=857 genes) and other functional classes (n=541 genes)^2^. This analysis demonstrated that cross-validation accuracy using TFs was comparable to all expressed genes for discriminating between 338 Subclasses^2^. (In the WMB taxonomy, the Subclass is considered an intermediate level of the atlas^2^ and comparable to the Group level in the HMBA dataset^29^.)

In the HMBA context, larger gene sets, such as the UniprotKB^49^ synapse related genes from (838 genes) and AnimalTFDB4^50^ transcription factors (1,119 genes) showed similar performance to the 4096 highly variable genes and outperformed the smaller set of extracellular matrix genes (414 genes) (Figure S1A). Surprisingly, using 4,096 randomly selected genes also had comparable performance to using the top highly variable genes, with similar batch correction and slightly worse biological conservation (Figure S1B-C). These results suggest that the number of genes used for integration may be as influential as their functional annotations.

Moreover, feature selection remains an important consideration for cross-species integration, particularly in datasets with uneven taxon sampling. In the spermatogenesis dataset, we tested a lineage-balanced HVG strategy designed to reduce potential bias toward the densely sampled primate clade. Although this alternative strategy did not substantially alter the integrated structure in this dataset, it produced comparable results to the standard HVG approach, suggesting that PIANO is relatively robust to this feature-selection choice (Figure S6). Nonetheless, such strategies may be useful in future cross-species analyses involving stronger taxonomic imbalance, greater phylogenetic distances, or lineage-specific technical variation.

### Model parameter selection

PIANO had reliable performance with a default model architecture of 3 hidden layers, 256 hidden nodes, and 32 latent dimensions and default training parameters of 200 max epochs, learning rate of 0.0002, and no weight decay, and generally similar performance over 256 different combinations of hyperparameters each for 5 datasets (Table S2). For the smaller and simpler single-species datasets, the negative log-likelihood of the data had a smaller magnitude than the more complex brain datasets. As such, we found a minimum improvement threshold of 0.1 (for early stopping of model training) to be more effective for these simpler datasets than the default threshold of 1.0 for the other datasets. Alternatively, early stopping can be turned off to train for a fixed number of epochs (Figure S1D).

The most important hyperparameter to adjust is the max Kullback-Leibler divergence (KL) beta-annealing hyperparameter (see Methods), where a higher value results in stronger batch correction in exchange for reduced biological conservation. For most integrations, we used the default max beta of 0.25 at the max number of training epochs (default 200). For the simpler single species datasets (besides the noisy lung dataset), we used a stronger max beta of 0.5 to increase batch correction. For datasets with greater diversity of cell types, such as the whole mouse brain single-cell and spatial transcriptomics atlas, we lowered the max beta value to 0.0625 to avoid over-regularizing the latent space. For the cross-species spermatogenesis dataset, which had a large evolutionary complexity (11 species) and only 96,464 cells, we used a 0.01 as a more conservative max beta.

The adversarial training and counts correction improved the batch correction for most integrations with little impact on biological conservation of cell types. In some situations, such as the noisy lung dataset or the *in silico* perturbation of withholding the non-human primate D1 SPNs in the basal ganglia atlas for cross species (due to their transcriptomic similarity to closely related D2 SPNs), it is recommended to turn off adversarial training or to not perform counts correction. Likewise, it is recommended to first identify cell types within each species before cross species integration to ensure signals from each species are still preserved.

### Limitations of this study

One limitation of the PIANO framework is that it relies on GPU acceleration for optimal runtime performance. Although the method can run on both x86 and ARM CPU architectures, its fastest runtimes rely on code compilation, which requires the use of a GPU that can support torch.compile optimizations. Moreover, these compilations introduce up to two minutes of additional compilation time overhead costs, which make up a larger proportion of runtime for smaller datasets. These hardware requirements can be met by most high-performance computing clusters (HPCs) or cloud computing platforms that support modern GPUs.

One limitation of the differentially expressed gene (DEG) analysis for the primate basal ganglia data is the small number of donors per species (i.e., only 4 marmosets). This would limit the power of bulk sequencing approaches for analyzing DEGs. On the other hand, each species has a large number of cells, which inflates the power of the Wilcoxon rank sums approach that was used to perform the DEG analyses and makes the p-value less informative. Instead, a stringent minimum gene expression proportion threshold of at least 10 counts per 100,000 transcripts in either comparison group was used to filter out genes with low expression, which helps avoid false positives.

One limitation of generative modeling for gene expression counts is the potential to over-correct for batch effects. For example, since PIANO uses a NB-GLM that assumes linear weights for the species covariate, naively regressing out all species effects may over-correct for species effects in a cell type that is not shared across all species. These issues can be minimized or avoided by only correcting for the batches that contain conserved cell types. Moreover, the generative modeling approach only considers the first order effects of perturbations on gene expression. For example, PIANO may not capture downstream effects after initial perturbations that may result from negative feedback loops or other gene-gene interactions. As such, PIANO’s generative modeling approach should be carefully utilized in conjunction with more comprehensive gene regulatory network analyses. Complex drug-drug interactions or non-linear disease patterns (e.g., an increase followed by a decrease) may benefit from more comprehensive modeling approaches.

### Related work

The negative binomial generalized linear model (NB-GLM) modeling used in the VAE decoder was inspired by the NB-GLM used by scTransform for normalizing scRNAseq data^51^. Other commonly used integration tools include Seurat (CCA)^8^, Harmony^7^, scVI^24^, and others. Additionally, some integration methods attempt to use a semi-supervised approach, such as scANVI^52^. However, these methods rely on the robustness of the assigned cell type labels and may bias results against novel biology or rarer cell types and may not model nuanced relationships between similar cell types. Moreover, semi-supervised training approaches may also introduce training instability (e.g., due to rare cell types) and longer runtimes.

Like PIANO, scTransform is a method for data normalization that can be utilized for differential gene expression (DGE) analyses^51^. The NB-GLM used in PIANO can be considered a more customizable data normalization approach that not only models sequencing depth but also enables customizable counts correction for multiple covariates. In contrast, methods such as DESeq2 are designed primarily for bulk RNA-seq data^53^. Although they can still be utilized for single-cell data via pseudobulking, these methods rely more heavily on the number of donors for statistical power. Rather than exclusively selecting one method for DGE analysis, utilizing multiple complementary approaches is recommended for establishing the robustness of results.

Some methods for integrating spatial transcriptomics data, such as SpaMask^54^, SpaCross^55^, SpaBatch^56^, and STAligner^57^, incorporate the spatial neighborhoods across cells or spots using graph neural networks. These methods can be used to align multiple adjacent slices of tissue but may struggle with computational scaling to larger datasets or datasets with disjoint or non-consecutive slices (i.e., aligning coronal and sagittal slices of brain tissue). In contrast, PIANO focuses on latent-variable modeling of the RNA gene counts in single-cell and spatial data, relying on the spatial coordinates of existing spatial slices. As such, PIANO can be used synergistically with these existing spatial integration methods. Future work may incorporate PIANO with similar graph attention mechanisms to expand spatial integration applications.

## Supporting information

Key Resources Table.

## Acknowledgements

This publication was supported by and coordinated through the Brain Initiative Cell Atlas Network (BICAN). This work was supported by National Institutes of Health UM1MH130981, DP2MH140136, and the Klingenstein–Simons Fellowship (F.M.K.). N.W. was supported by The McDonnell Fellows in Neuroscience at Princeton University and National Institutes of Health 5T32MH065214. This work was completed in part at the Princeton Open Hackathon, part of the Open Hackathons program. The authors would like to acknowledge OpenACC-Standard.org for their support. The authors would like to acknowledge Dr. Milos Nikolic for his writing advice during the preparation of this manuscript. The authors would like to acknowledge the Allen Institute for Neural Dynamics for their computational support with Code Ocean. We wish to thank the Allen Institute founder, Paul G. Allen, for his vision, encouragement, and support.

## METHODS

### PIANO model architecture

PIANO uses approximate Bayesian inference to train a modified variational autoencoder (VAE)^9^. For the observed gene counts X, the recognition model (encoder) uses a fully-connected feed-forward deep neural network (DNN) to learn a latent variable 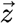. The latent variable is a diagonal Gaussian, with independent means and standard deviations for each latent dimension.

The generative model (decoder) uses the latent variable concatenated with the covariates as inputs to a DNN (mirroring the encoder), which reconstructs each normalized gene expression proportion *x*_*gene,prop*_ using the softmax transformation, where 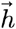 is the DNN output:

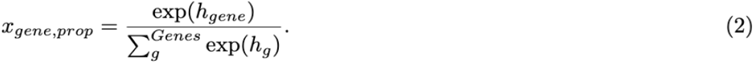

Reflecting the discrete nature of mRNA transcripts^58^, the generative model learns a negative binomial (NB) distribution for each gene:

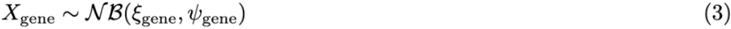

A generalized linear model (GLM) is used to perform regression over the covariates. The NB distribution for each gene is parameterized by ξ_*gene*_ (Ksi) and ψ_*gene*_ (Psi)^59^, where:

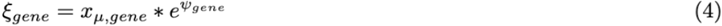

And:

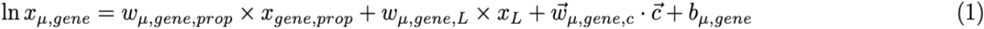

Here, *x*_*gene,prop*_ is the proportion expressed for a gene in a given cell. This proportion is scaled by the library size *x*_*L*_(total number of transcripts) for each cell, which influences the mean gene expression^51^. The vector of the covariates 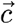 includes categorical covariates (one-hot encoded) and continuous covariates (z-scored), 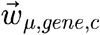 is a vector of weights for the covariates, 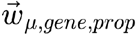 is a scalar weight for the gene proportion,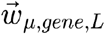 is a scalar weight for the library size, and the bias term is given by *b*_*μ*_,_*gene*_. This differs from scVI’s implementation, which does not include a bias term and uses weights for only the primary batch key^24^ as opposed to multiple covariates:

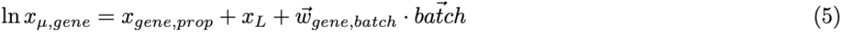

PIANO also models the other NB parameter:

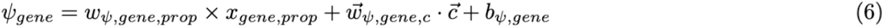

Here,*w*_*ψ*_,_*gene,prop*_ is the scalar weight for the gene proportion, *w*_*ψ*_,_*gene*_,_*c*_ is the vector of weights for the covariates, and *b*_*ψ*_,_*gene*_ is the bias term. We assume that the library size should only impact the mean expression of genes, and therefore, do not include it to model the log-odds. scVI uses a mean-overdispersion parameterization^24^, which does not model log-odds.

The likelihood of the data under the PIANO NB-GLM is given by:

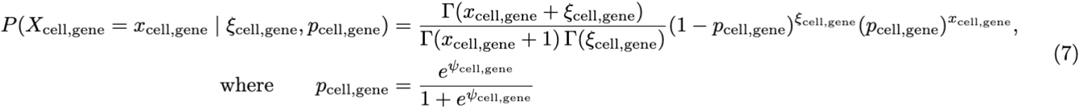

### PIANO evidence lower bound objective

Using the reparameterization trick^9^, PIANO is trained by minimizing an evidence lower bound objective (ELBO), which is a lower bound on the log-probability of the observed gene expression data given the covariates. The ELBO is the sum of a data fidelity term, given by the average log-likelihood of the data under the NB observation model, and the negative Kullback-Leibler divergence (KL) between the approximate posterior over Z from the recognition model and the isotropic Gaussian prior. The total ELBO is the sum of ELBOs across cells. The ELBO can be derived as follows, where are the parameters of the decoder *ϕ* and are the parameters of the encoder:

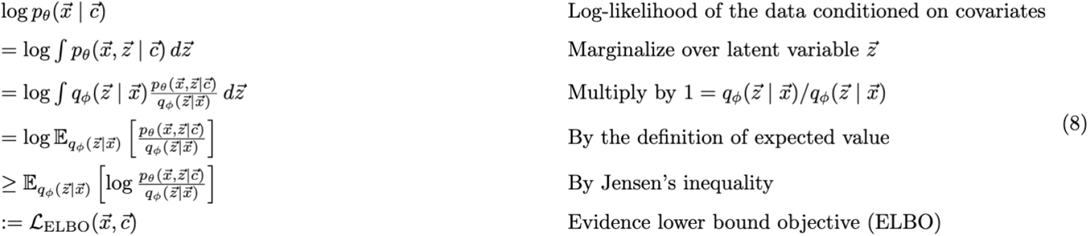

The ELBO can be written as an expected log-likelihood and a KL term^9^. For a given cell, we have:

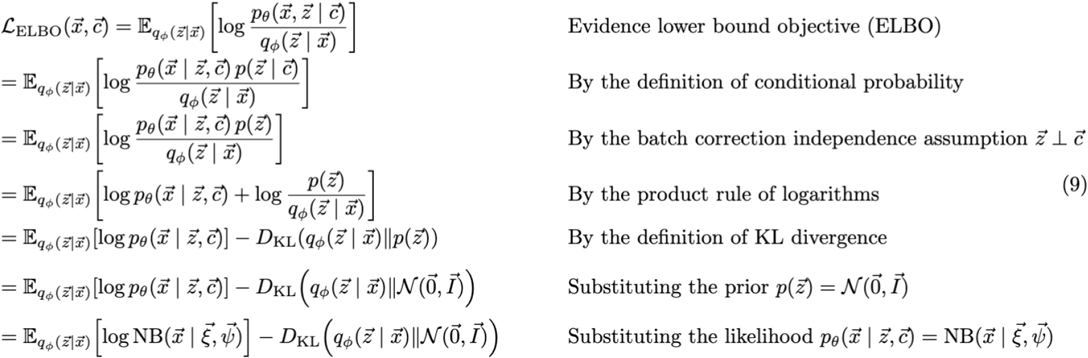

Here, we have:

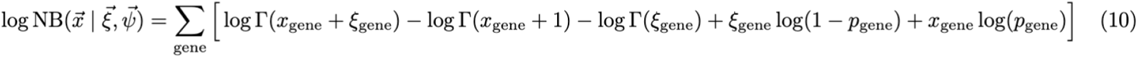

where 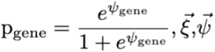 are the NB-GLM parameters from the generative model using 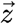 and 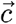 as inputs, and is the Gamma function.

The KL term, which involves the latent variable (a diagonal Gaussian) and an isotropic Gaussian prior distribution, can be calculated analytically^9^. The term 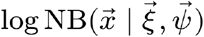 is evaluated using the log-likelihood of the original counts under the NB-GLM, with the *ξ*_*gene*_,*ψ*_*gene*_ terms obtained from the forward pass of the VAE. By sampling from the latent variable, the generative model produces the expectation over the log-likelihood of the data. This VAE model architecture enables PIANO to train in an unsupervised manner using backpropagation^60^.

### Gradient reversal layer

PIANO enables adversarial training via gradient reversal^10^. This is implemented using an adversarial network that utilizes the latent variable as its input and predicts the corresponding categorical covariates (i.e., batches). This adversarial network is trained to minimize the binary cross-entropy (BCE) per covariate. A gradient reversal layer placed between the encoder and the adversarial network flips the sign of the gradients during backpropagation^10^, which allows PIANO to simultaneously train the adversarial network to improve at predicting the categorical covariates from the latent space 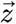 whereas the encoder is trained to make the latent space less informative of these categorical covariates 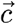. Specifically, the model aims to minimize the loss, where ρ are the parameters of the adversarial network:

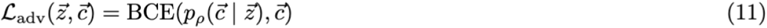

### Implementation details

The encoder, or recognition model, learns an approximate posterior distribution over the latent P(Z|X) (e.g., 32 latent dimensions) given a vector of gene counts (e.g., 4096 highly variable genes); specifically, the encoder outputs a probabilistic posterior latent representation for each cell using an independent mean and standard deviation for each latent dimension.

Using the reparameterization trick^9^, a z-score is sampled outside of the computational graph to produce the posterior latent representation P(Z|X) from the mean, standard deviation, and z-score. This latent representation is concatenated with covariates as the input to the decoder.

Given a latent vector and a set of covariates, C, (e.g., one-hot encoding for 3 primate species produces 3 dimensions to concatenate with the latent vector), the decoder parametrizes P(X|Z,C) (i.e., a vector for 4096 genes), which is the normalized gene expression; specifically, the decoder uses the softmax function to obtain gene proportions that sum to one. These proportions are then scaled by the library size (total counts) of a given cell to obtain the means of the negative binomial distributions over the selected genes. Compressing the data through the VAE bottleneck helps the latent space representation capture only the most important biological information for reconstructing the gene proportions. Feeding the covariates into only the decoder helps the model remove batch effects from the latent space while still modeling the original gene counts, which requires information from the covariates.

Both the encoder and decoder are implemented in PyTorch using a multi-layer perceptron (MLP), or feedforward neural network, following the pattern of feed-forward linear layer, batch normalization, rectified linear unit (ReLU), and dropout. PIANO uses a default, symmetric model architecture with 3 layers, 256 hidden nodes per linear layer, and 32 latent dimensions for the encoder and decoder. PIANO supports the option of utilizing adversarial training. The gradient reversal layer is implemented by a sign change during the backwards step during backpropagation^10^. This layer is placed between the encoder and adversarial network, which is implemented using a 2 layer network that uses the same dimensions as the latent space as input, same number of hidden nodes as the encoder and decoder, a ReLU activation layer, and the same number of dimensions for output as the number of categorical covariates.

PIANO uses default parameters with 1e-5 for the batch normalization epsilon, 1e-1 for batchnorm momentum, and 0.1 for dropout rate. PIANO uses an AdamW optimizer with a default learning rate of 2e-4, mini-batch size of 128 cells, and no weight decay. For training stability, datasets with normalized total transcript counts in the hundreds of thousands per cell are first mean-scaled to 10,000 transcripts per cell within each batch; the gene expression data are also log1p-transformed before being used as input for the encoder. For each training step, we use a mini-batch update with 128 randomly sampled cells without replacement.

### Beta annealing for Kullback-Leibler divergence

Following the Beta-VAE approach, the most important PIANO hyperparameter is the Kullback-Leibler (KL) divergence beta parameter, which is an additional term that acts as a weight on the KL for the ELBO calculation^61^. This beta parameter controls the regularization of the latent space, which serves to govern the tradeoff between biological conservation and batch correction. Specifically, we have:

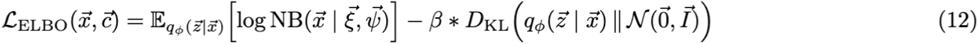

To mitigate KL vanishing, VAEs are trained using beta-annealing^62^, starting with a small beta value and increasing as the model trains. PIANO uses a linear beta-annealing schedule that starts from an initial value of 0.0, increases linearly after every training step, and reaches the max beta value after n_annealing_epochs, which has a default value of 0.25. By default, n_annealing_epochs is set to the max number of training epochs, which has a default value of 200 epochs. PIANO finishes training after the max number of training epochs unless early stopping is activated. This occurs if the total sum of the raw losses (negative log-likelihood, KL, and adversarial loss) no longer improves by the min_delta (default 1.0) after several epochs (a default patience of 5 epochs). The KL beta hyperparameter is not used for the early stopping calculation to avoid conflicts with its linearly-increasing annealing schedule. PIANO trains to minimize the overall loss:

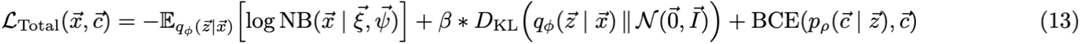

### Performance improvements

PIANO’s speed and scalability improvements are driven by its efficient implementation. PIANO is implemented in vanilla PyTorch and achieves faster training through backpropagation compared to existing deep learning methods through better utilization of GPU code compilation. Using torch.compile, PIANO compiles the entire backpropagation training loop, including both forward and backward passes. Utilizing a custom, lightweight code implementation, PIANO minimizes graph breaks in the computation graph, which enables better GPU utilization. These improvements are possible through numerous code optimizations, such as avoiding expensive PyTorch lightning callback functions (i.e., by instead using custom early-stopping monitors) and designing simplified control logic (i.e., avoiding expensive code branch predictions).

Moreover, PIANO uses custom PyTorch Dataloaders and Dataset classes to ensure scalability to truly large datasets. As a deep learning model trained with backpropagation using mini-batch stochastic gradient descent, PIANO supports training on multiple separate AnnData objects without concatenation. This enables sampling cells from multiple objects for the same mini-batch update as if the data were stored in the same object with minimal overhead costs. Additionally, PIANO supports multiple memory modes to maximize speed improvements. The dense GPU mode keeps the entire data in dense GPU memory for the absolute fastest training speeds, the custom sparse GPU mode enables storing data in sparse GPU tensors, the dense CPU mode enables balanced performance between speed and memory, and the sparse CPU mode offers the most memory efficient implementation with similar speed to the other modes. Furthermore, this mode also supports sparse continuous covariates, where PIANO dynamically processes the metadata without needing to create a dense covariates matrix. This further reduces the memory costs for learning comprehensive models on large atlases.

### Preprocessing

Publicly available and preprocessed datasets were used in all integrations; as such, no new quality control thresholds were used to remove low quality cells. Feature selection is an important consideration for data integration. For cross-species comparisons, the data were subset to the human one-to-one orthologs from Ensemble version 114^63^. For the primate basal ganglia integration gene sets, one-to-one ortholog transcription factors were selected from AnimalTFDB4^50^ and one-to-one ortholog functional gene sets were queried from UniProtKB^49,64^. An implementation of Scanpy’s sc.pp.highly_variable_genes was used to select 4096 genes for most integrations, using the ‘seurat_v3’ flavor and the most prominent source of variation for the batch key^31,41^. A hyperparameter sweep was performed over the number of top HVGs, and using 4096 HVGs for integrating the HMBA data had robust performance (Figure S3A).

For the single species dataset integrations, the batch keys used were sequencing platform for pancreas, donor for lung, dataset batch for immune, donor for tabula muris, and donor for heart atlas. For each of the cross-species integrations, species was used as the batch key. For the whole mouse brain multi-modal integration, all 1,122 genes were used. For the whole mouse brain spatial integration, all 477 overlapping genes were used. For the spermatogenesis dataset, 3,048 one-to-one orthologs were identified across the 11 species, and 2,000 highly variable genes were selected with species as the batch key for integration and pseudotime analysis. For this dataset, an alternate strategy was explored for selecting features for cross-species integration to reduce bias towards densely sampled clades. The top 2,000 highly variable genes (HVGs) were first identified independently within each species using the Seurat v3 flavor in Scanpy^31^. Species-specific HVG lists were then aggregated into broader phylogenetic groups (primates, mouse, opossum, platypus, and chicken), and genes were ranked within each group by their normalized species support, median HVG rank, and best HVG rank. The final set of 2,000 genes was selected by round-robin sampling across groups, yielding a balanced feature set that represented both densely and sparsely sampled lineages. For the SEA-AD dataset, features were selected by union rather than by HVG selection alone. The top 4,096 HVG’s from the snRNA-seq data were first identified using the Seurat v3 flavor in Scanpy. To enable joint modeling with MERFISH data, MERFISH panel genes not already among the selected HVGs (37) were included, and this set was further supplemented with disease-relevant genes (118). The final feature set contained 4,251 genes, which were used for training. For the Tahoe-100M dataset, the top 4096 HVGs were selected with plates as the batch key.

For most datasets, the raw gene counts were used as input. For datasets with substantially larger magnitudes for the raw data (i.e. due to normalizing differences in the sequencing data acquisition), each dataset was scaled to a mean of 10k. These datasets include ‘Villani’ for the immune integration, the ‘fluidigmc1’, ‘smarter’, and ‘smartseq2’ datasets for the pancreas integration, and all donors for the tabula muris integration.

To balance the number of cells between datasets, 6400 cells per group were used for each species in the primate basal ganglia, 6,400 cells per Subclass for each modality (scRNAseq, spatial) in the whole mouse brain integration, and 2^16^ (65,536) cells per developmental timepoint for each species (mouse and marmoset) for the developmental integration. For the Tahoe-100M dataset, all cells were used for training, with a subset of cells used for visualization.

For principal component analysis (PCA), 30 principal components were used in each integration example. For UMAP visualization, the Scanpy^31^ default 15 neighbors were used to compute the neighborhood graph. For Seurat, the k.weight was set to 50 (as opposed to the default 100 to enable integrations on batches with fewer cells). For Harmony, default parameters were used (with max 20 iterations). For scVI, the dispersion for its zero-inflated negative binomial distribution (as described in their original manuscript) was set to ‘gene-batch’. All other parameters were set to defaults besides the model architecture, which was set to use the same 3 hidden layers, 256 hidden nodes per hidden layer, and 32 latent dimensions used by PIANO (and the other deep learning methods) for benchmarking comparisons.

### Benchmarking

Integration embeddings were evaluated by biological conservation metrics, which measure separation of cell type clusters, and batch correction metrics, which measure how mixed cells are between different batches. The biological conservation metrics used were normalized mutual information (NMI) and adjusted Rand index (ARI). The batch correction metrics used were the k-nearest-neighbors batch effect test (KBET) and inverse local Simpson’s index (iLISI)^27^. It is well understood that a good integration should maintain separation of cell type clusters while mixing data between batches without over correction^27^.

The scib_metrics package was used to run integration benchmarking with commonly used metrics^27^ The average of the Leiden Adjusted Rand Index (ARI) and Leiden Normalized Mutual Information (NMI) scores was used to measure biological conservation; both metrics use de novo Leiden clustering on the latent space representation of the data and compare the clusters to the ground truth labels^27^. The average of the K-nearest neighbors Batch Effect Test (KBET) and integration Local Inverse Simpson’s Index (iLISI) scores were used to quantify batch correction; these metrics compare the number of cells from different batches in the neighborhood of a given cell^27^.

### Hyperparameter tuning

PIANO was run using the default architecture and parameters unless otherwise noted. For the spermatogenesis dataset, alternative network sizes were evaluated and KL values to assess the effect of model hyperparameters on integration performance (Figure S6). The final model used a reduced network architecture with 64 nodes per layer and a KL beta value of 0.01 (Figure 4, S6). PIANO with the default architecture was also used as a comparison (Figure S6). To assess the effect of feature selection, the standard highly variable gene selection strategy was compared with the lineage-balanced approach (described above) that was designed to reduce overrepresentation of the densely sampled primate clade (Figure S6).

### Differential Gene Expression

Two categories of differentially expressed genes (DEGs) were computed. In the first category, for each Subclass level cell type, comparisons were made between each non-human primate and humans. A comparison was performed between the DEGs derived from raw gene expression counts and the PIANO (donor and species) corrected counts. Additionally, global DEGs were computed between species (using cells from all cell types) using the raw counts. For each Subclass, set differences between DEGs obtained using raw and corrected counts were compared to the global species DEGs to quantify genes that were not Subclass specific.

In the second category, pairwise comparisons were performed between all pairs of Subclasses within the same species. The genes with higher expression in a Subclass compared to all other Subclasses were treated as Subclass DEGs for a given species, and the intersection of these DEGs were computed across species to obtain conserved Subclass DEGs. The DEGs were computed using the raw counts and the PIANO (same-species donor) corrected counts.

The Scanpy Wilcoxon Rank-Sum test^31^ was used for each DEG comparison, with default parameters besides specifying the comparison groups. A minimum (base 2) log-fold change of 0.1 and adjusted p-value less than 0.01 was required to determine statistical significance. To reduce false positives, the DEG comparison group with the higher expression was required to have a minimum expression proportion of at least 10 raw counts per 100,000 unique transcripts.

### Multi-species developmental pseudotime analysis

Pseudotime trajectories were calculated using diffusion pseudotime from Scanpy^31^. A cell-cell neighborhood graph was first constructed from the PIANO latent space using 50 nearest neighbors and a Gaussian kernel. Diffusion maps were calculated from this graph with 10 diffusion components. The trajectory was rooted in the undifferentiated spermatogonial population, based on prior cell-type annotations. Specifically, the centroid of all cells annotated as undifferentiated spermatogonia was calculated from the PIANO latent space, and the cell closest to this centroid was selected as the root cell. Diffusion pseudotime was then computed with the function sc.tl.dpt using the first 10 diffusion components.

### Multi-modal modeling of Alzheimer’s disease progression

For PIANO’s integration of the snRNA-seq data, highly variable genes (HVGs) were selected using Scanpy’s sc.pp.highly_variable_genes with the ‘seurat_v3’ flavor (4096 genes)^31^. The spatial genes from the MERFISH panel (37 genes) which were not present in the top 4,096 HVGs and gene markers related to Alzheimer’s disease (118) were also included, which brought the total number of genes used for the single-nucleus integration to 4,251 genes.

For spatial coordinate assignment, scRNA-seq and spatial dataset were separated by class (Glutamatergic, GABAergic, and non-neuronal) and integrated across modalities with PIANO (default parameters, 140 genes used). Within PIANO’s latent space, k-nearest neighbor (KNN) transfer (k=1) was used to assign each snRNA-seq cell the spatial coordinates of its closest MERFISH neighbor. Each snRNA-seq cell was assigned a transferred subclass label. A strong diagonal enrichment in the confusion matrix between assigned scRNA-seq subclass and KNN transferred subclass confirms accurate spatial coordinate assignment (Figure S5A).

For the generative model, donor ID and sex were included as categorical covariates, while age at death (Z-scored) and CPS were modeled as continuous covariates. CPS values were taken directly from the SEA-AD donor-level annotations and were not recomputed. Donor ID was used as the batch key, and all other parameters were set to defaults. Generated expression was retrieved at eleven fixed CPS values spanning 0 to 1 in increments of 0.1. Generated and observed counts were normalized to counts per 10,000 and log1p transformed.

After generating expression across disease progression, the assigned spatial coordinates of scRNA-seq cells were plotted on a spatial section. A single representative MERFISH section (H21.33.031.CX24.MTG.02.007.1.01.01) was selected due to its clearly resolved cortical layers. The intact upper piece (with spatial y-coordinate > 3,500) was selected for improved visualization. All MERFISH cells from this section were plotted in grey to outline the tissue and its cortical layers. The snRNA-seq cells of the subclass of interest, positioned at their KNN-transferred coordinates, were then plotted over this outline and colored by generated expression at each CPS value. For each cell, generated counts were normalized to counts per 10,000 and then log1p transformed. To emphasize disease-related changes, expression was displayed as the difference relative to baseline for Early (CPS value of 0.2), Middle (0.5), and Late (1.0) stages of disease (Δ = expression at a given CPS minus expression at CPS = 0) for *APOE, MAPK8, MME*, and *APP*.

For the SST Supertype vulnerable and unaffected group analysis, each snRNA-seq cell was assigned the Supertype of its nearest MERFISH neighbor and labeled vulnerable or unaffected if that transferred Supertype belonged to the corresponding group, with all other cells left unlabeled (S5B). The vulnerable group consists of SST_2, SST_11, SST_19, SST_20, SST_22, SST_23, and SST_25, and the unaffected group consists of SST_1, SST_4, SST_5, SST_7, SST_9, SST_10, and SST_13. The same transferred Supertype labels were used to resolve *APP* expression within the L2/3 IT subclass (S5C). Cells were subset to the L2/3 IT supertypes occupying the section (L2/3 IT_1 and L2/3 IT_6 in L2; L2/3 IT_10 and L2/3 IT_13 in L3) and plotted separately at CPS values of 0 (Before Onset), and the expression relative to baseline at values of 0.5, and 1.0 (S5D).

### Generative modeling of cancer drug perturbations

A generative model was learned over all cells. For each simulated perturbation, the data was subset to the cell line and drug of interest for all 3 non-zero dosages and the control group. This approach enables specifying the cell line to model. Using these subset cells, an integrated latent representation was sampled (using the model trained over all cells), and generative modeling was performed (again, using the model that was trained over all cells) to simulate gene expression at each drug concentration, marginalizing across all plates.

## Code availability

The PIANO source code is publicly available at https://github.com/NingWang1729/piano with an installable package at https://pypi.org/project/piano-integration.

## Data availability

All datasets used were already preprocessed and publicly available. Download links can be found in the key resources table.

## Author Contributions

N.W., C.L., J.W.P., and F.M.K. designed the method. N.W. and D.T. implemented the method. N.W., C.C., H.F., and V.N. benchmarked other methods. N.W., C.C., V.N., H.F., N.S., C.L., D.Y., Z.Y., C.C., M.D., S.D., L.C., and J.S. performed data collection, preprocessing, and analysis. N.W., C.C., V.N., C.L., J.W.P., and F.M.K. wrote the manuscript with input from all authors.

## Supplemental information

Document S1. Figures S1–S6.

Tables S1–S6. Key Resources Table.

