## Supplementary material for "PIANO: Probabilistic Inference Autoencoder Networks for multi-Omics enables robust generative modeling of gene expression and scales single-cell integration to 100 million cells": Document S1. Figures S1-S6.

Supplemental Information Index:

- Figure S1, related to Figure 2: Primate basal ganglia integration.
- Figure S2, related to Figure 2: *In silico* perturbations of primate basal ganglia cell types.
- Figure S3, related to Figure 2: Subclass-specific differential gene expression comparisons across species.
- Figure S4, related to Figure 2: Evolutionarily-conserved Subclass-defining differentially expressed genes
- Figure S5, related to Figure 6: Validation of spatial coordinate transfer and layer-specific gene expression across disease progression
- Figure S6, related to Figure 4: Spermatogenesis integration across evolutionarily distant species
- Table S1, related to Figure 1: Integration benchmarking on single-species datasets
- Table S2, related to Figure 1: PIANO performance across hyperparameters
- Table S3, related to Figure 2: Primate basal ganglia integration
- Table S4, related to Figure 2: Model architecture comparisons
- Table S5, related to Figure 3: Cross-species developmental brain integration
- Table S6, related to Figure 5: Whole mouse brain integration
- Key Resources Table

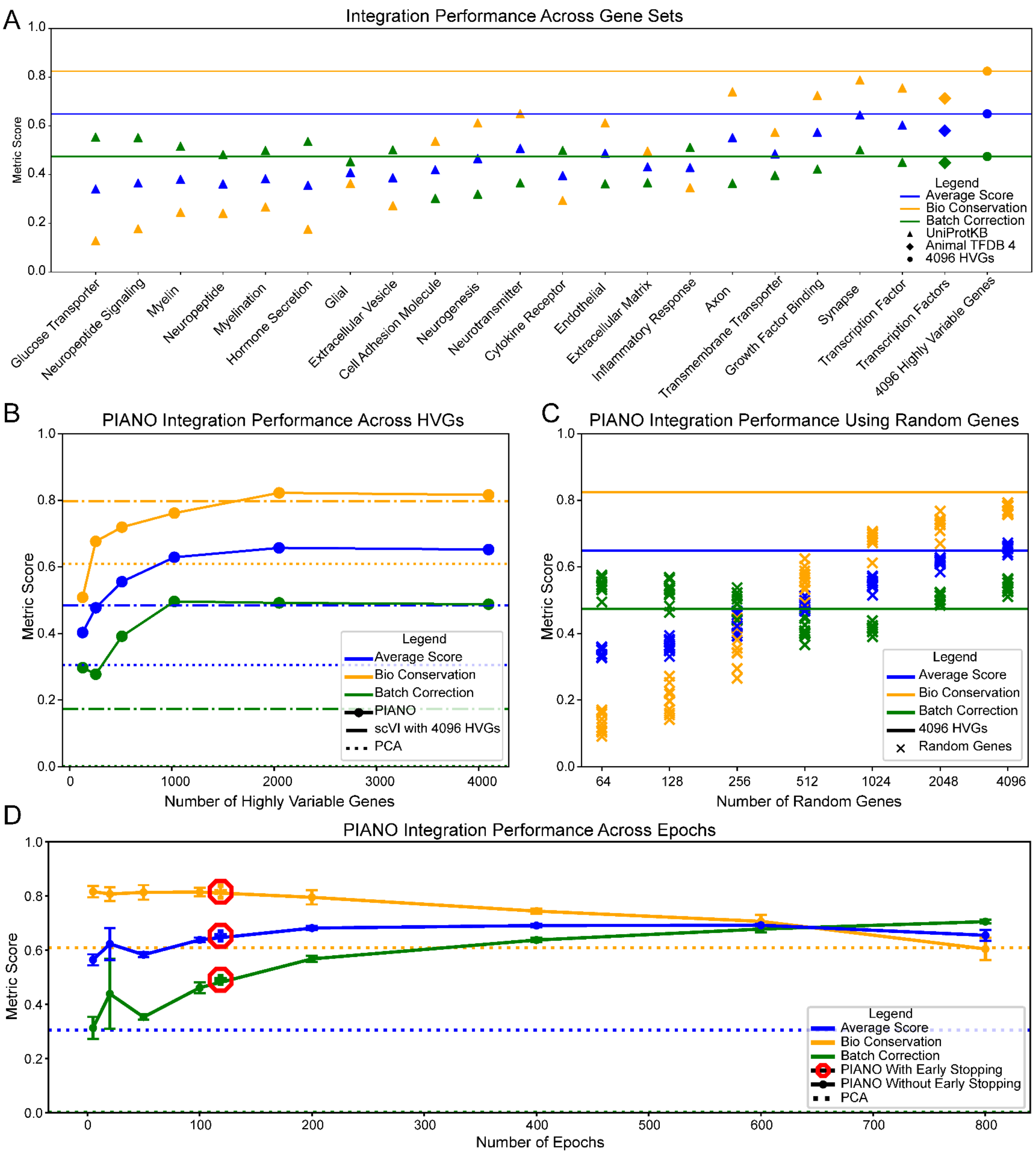


Figure S1, related to Figure 2: Primate basal ganglia integration. A) Functional gene sets show the influence of gene set size on integration performance. B) Using more highly variable genes improves integration performance, with similar performance at 2048 and 4096 highly variable genes. C) Similarly, large numbers of randomly selected genes gives improved integration performance but has lower biological conservation than using highly variable genes. D) Early stopping shows robust performance, with a larger number of training epochs leading to stronger batch correction at the expense of biological conservation.

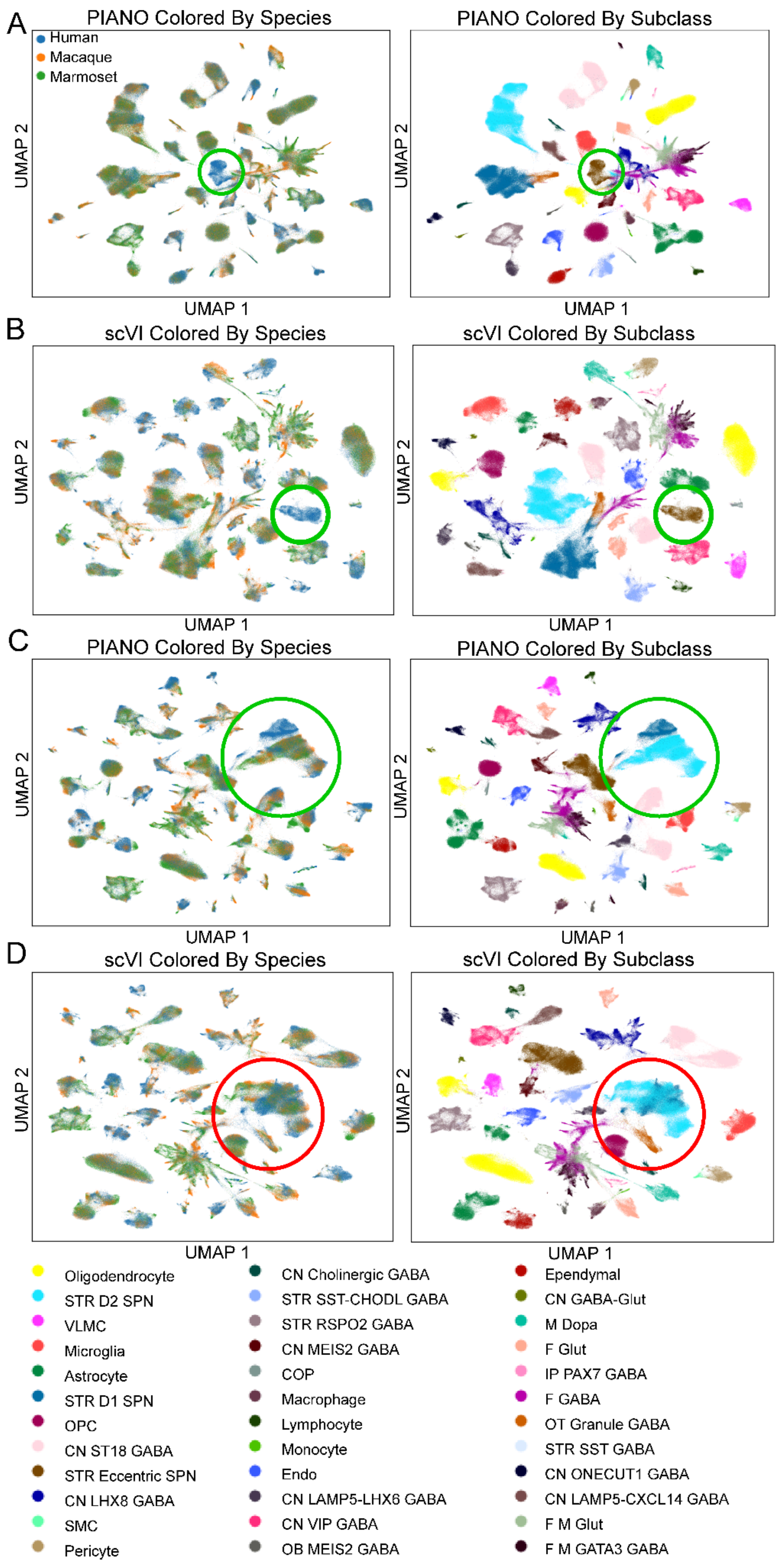

A
PIANO Colored By Species
Human
Macaque
Marmoset
UMAP 2
UMAP 1
PIANO Colored By Subclass
UMAP 2
UMAP 1
B
scVI Colored By Species
UMAP 2
UMAP 1
scVI Colored By Subclass
UMAP 2
UMAP 1
C
PIANO Colored By Species
UMAP 2
UMAP 1
PIANO Colored By Subclass
UMAP 2
UMAP 1
D
scVI Colored By Species
UMAP 2
UMAP 1
scVI Colored By Subclass
UMAP 2
UMAP 1
Oligodendrocyte
STR D2 SPN
VLMC
Microglia
Astrocyte
STR D1 SPN
OPC
CN ST18 GABA
STR Eccentric SPN
CN LHX8 GABA
SMC
Pericyte
CN Cholinergic GABA
STR SST-CHODL GABA
STR RSPO2 GABA
CN MEIS2 GABA
COP
Macrophage
Lymphocyte
Monocyte
Endo
CN LAMP5-LHX6 GABA
CN VIP GABA
OB MEIS2 GABA
Ependymal
CN GABA-Glut
M Dopa
F Glut
IP PAX7 GABA
F GABA
OT Granule GABA
STR SST GABA
CN ONECUT1 GABA
CN LAMP5-CXCL14 GABA
F M Glut
F M GATA3 GABA

Figure S2, related to Figure 2: *In silico* perturbations of primate basal ganglia cell types. A-B) PIANO and scVI capture the human-exclusive STR Eccentric SPN cell type after withholding the non-human primate cells. C-D) In the more difficult D1 MSN perturbation, PIANO preserves the simulated human-specific D1 MSN when not using adversarial training and species-correction whereas scVI loses human-specificity.

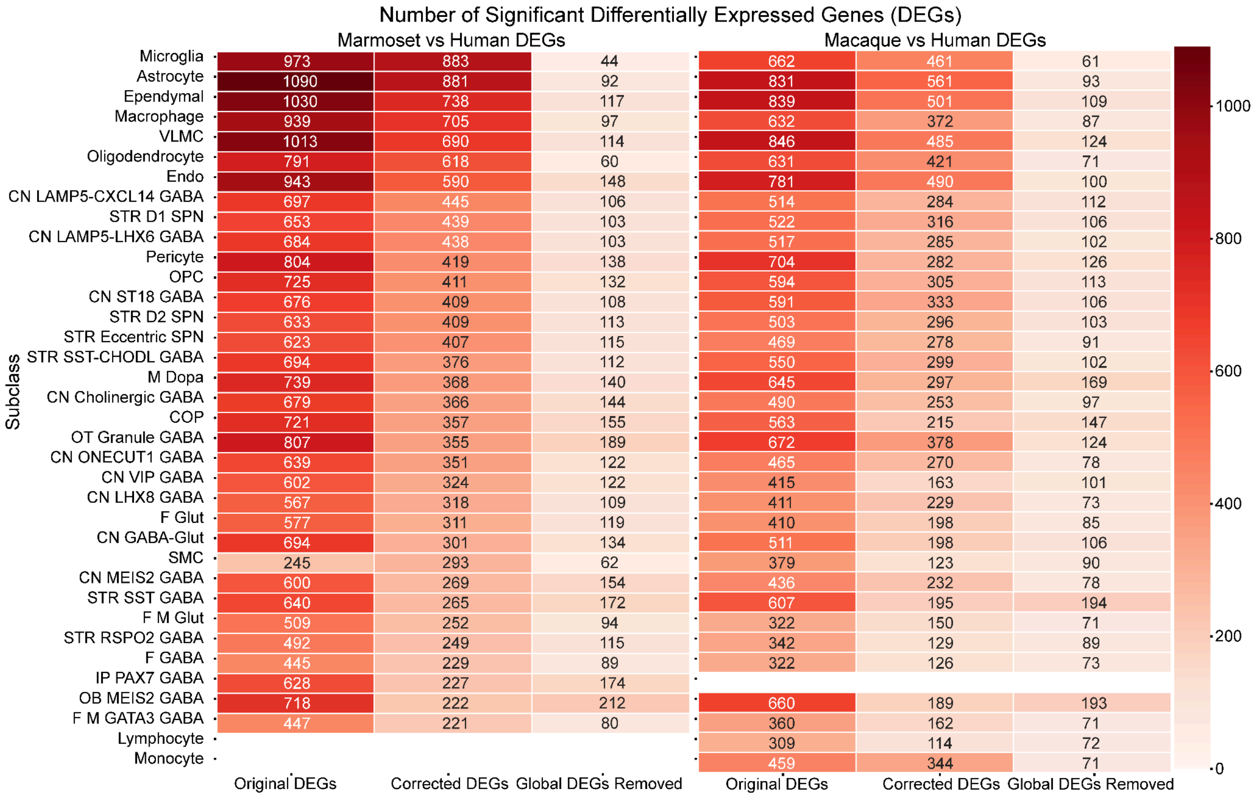


Figure S3, related to Figure 2: Subclass-specific differential gene expression comparisons across species. PIANO corrected counts produce more stringent differentially expressed genes (DEGs) compared to using raw counts, potentially avoiding false positives. This includes removing genes that are global DEGs as opposed to Subclass-specific. The results show that the non-neuronal cell types (at the Subclass resolution) generally have more DEGs compared to the neuronal cell types, suggesting greater evolutionary divergence.

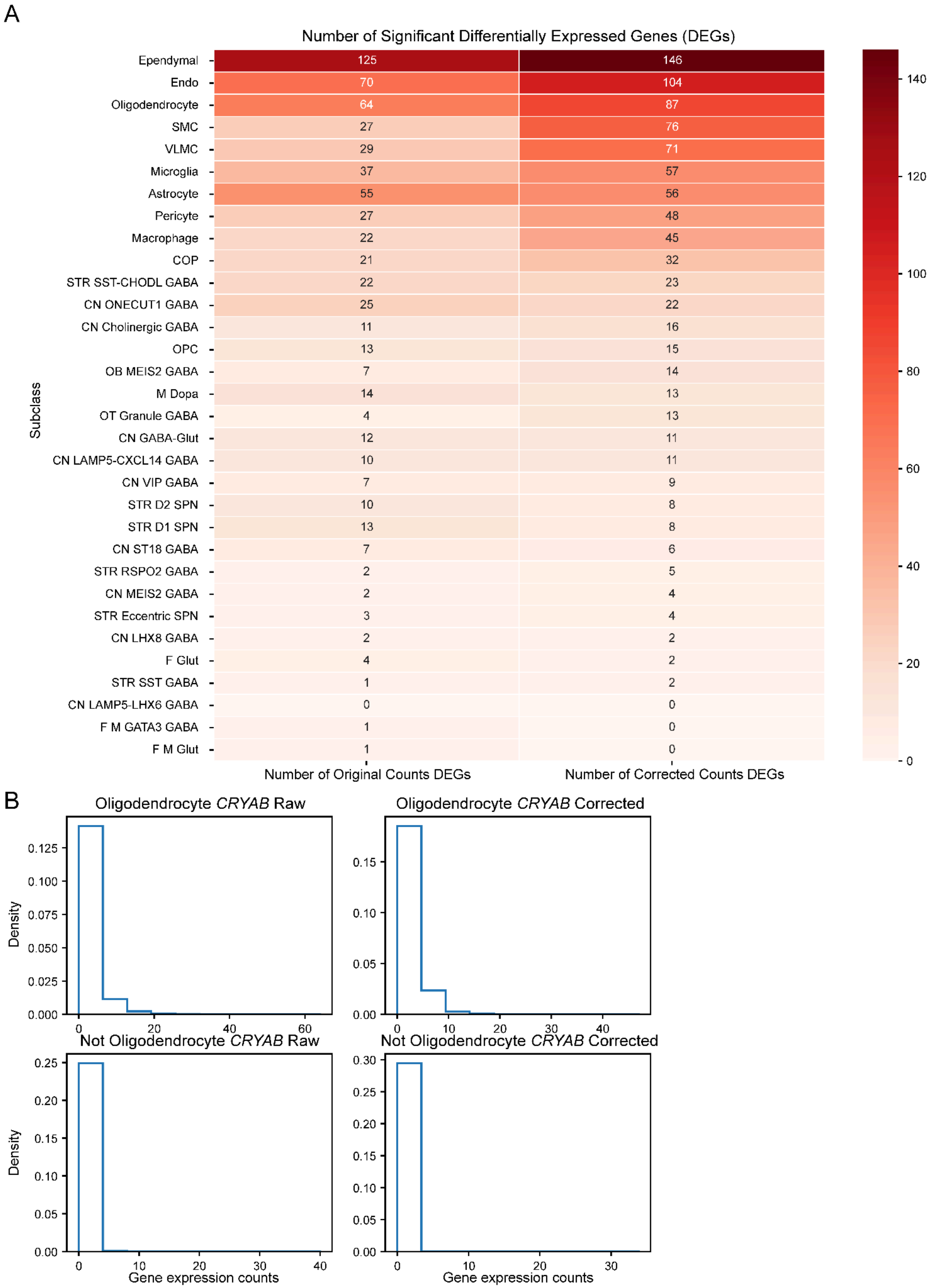
A
Number of Significant Differentially Expressed Genes (DEGs)
Subclass
Ependymal 125 146
Endo 70 104
Oligodendrocyte 64 87
SMC 27 76
VLMC 29 71
Microglia 37 57
Astrocyte 55 56
Pericyte 27 48
Macrophage 22 45
COP 21 32
STR SST-CHODL GABA 22 23
CN ONECUT1 GABA 25 22
CN Cholinergic GABA 11 16
OPC 13 15
OB MEIS2 GABA 7 14
M Dopa 14 13
OT Granule GABA 4 13
CN GABA-Glut 12 11
CN LAMP5-CXCL14 GABA 10 11
CN VIP GABA 7 9
STR D2 SPN 10 8
STR D1 SPN 13 8
CN ST18 GABA 7 6
STR RSPO2 GABA 2 5
CN MEIS2 GABA 2 4
STR Eccentric SPN 3 4
CN LHX8 GABA 2 2
F Glut 4 2
STR SST GABA 1 2
CN LAMP5-LHX6 GABA 0 0
F M GATA3 GABA 1 0
F M Glut 1 0
Number of Original Counts DEGs
Number of Corrected Counts DEGs
B
Oligodendrocyte *CRYAB* Raw
Oligodendrocyte *CRYAB* Corrected
Not Oligodendrocyte *CRYAB* Raw
Not Oligodendrocyte *CRYAB* Corrected
Density
Gene expression counts

Figure S4, related to Figure 2: Evolutionarily-conserved Subclass-defining differentially expressed genes. A) PIANO recovers DEGs that are conserved across species. B) The *CRYAB* gene is not identified as a Oligodendrocyte-defining DEG in Marmoset using raw counts due to outliers with large values in the non-Oligodendrocytes (which extend the x-axis to 40 counts). The corrected counts reduce the effects of outliers, indicated by the x-axis stretching to 30 counts, which help recapitulate *CRYAB* as a evolutionarily-conserved Oligodendrocyte DEG (as opposed to human specific).

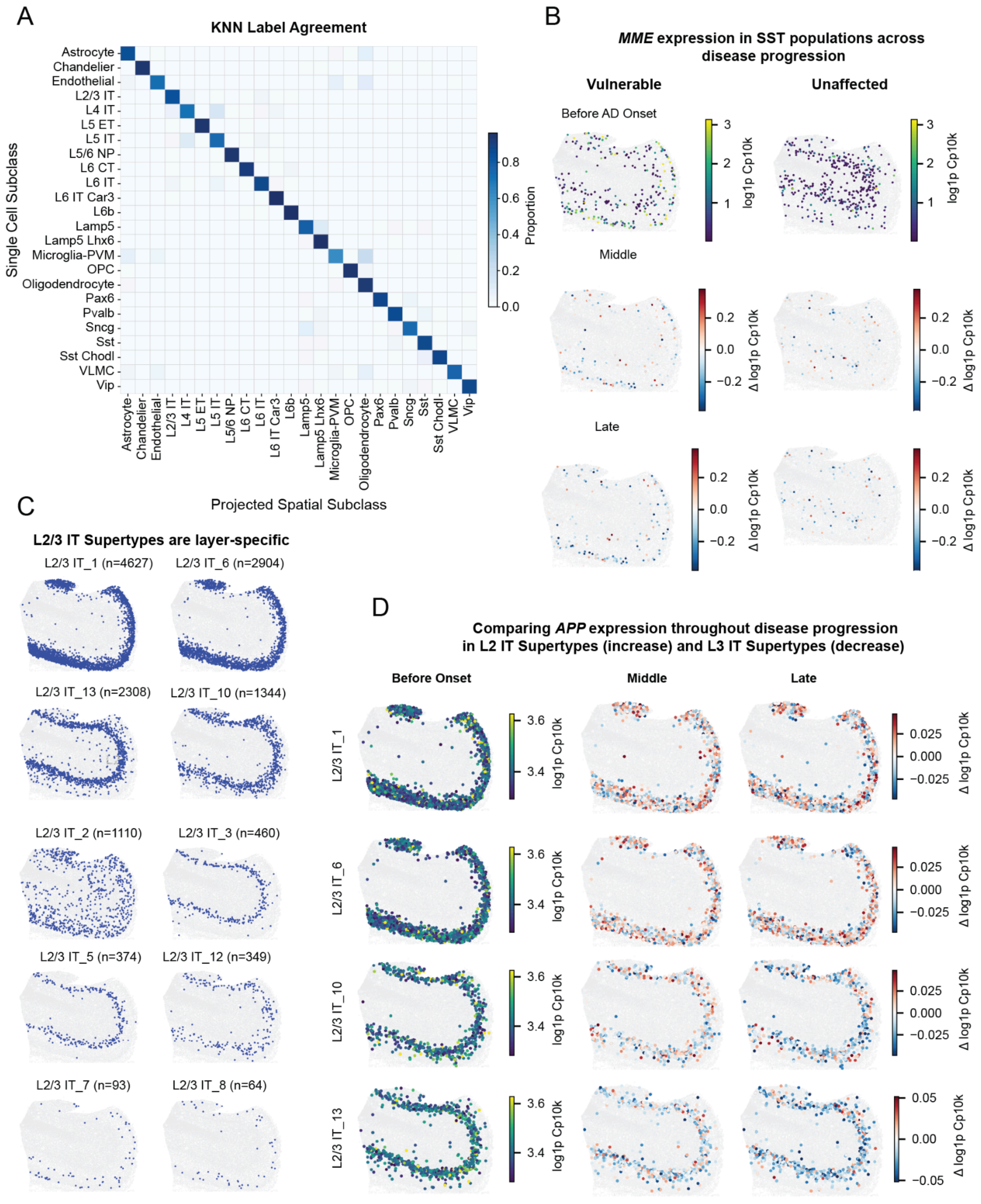

A
KNN Label Agreement
Single Cell Subclass
Projected Spatial Subclass
Proportion
Astrocyte
Chandelier
Endothelial
L2/3 IT
L4 IT
L5 ET
L5 IT
L5/6 NP
L6 CT
L6 IT
L6 IT Car3
L6b
Lamp5
Lamp5 Lhx6
Microglia-PVM
OPC
Oligodendrocyte
Pax6
Pvalb
Sncg
Sst
Sst Chodl
VLMC
Vip
B
MME expression in SST populations across disease progression
Vulnerable
Unaffected
Before AD Onset
Middle
Late
log1p Cp10k
Δ log1p Cp10k
C
L2/3 IT Supertypes are layer-specific
L2/3 IT_1 (n=4627)
L2/3 IT_6 (n=2904)
L2/3 IT_13 (n=2308)
L2/3 IT_10 (n=1344)
L2/3 IT_2 (n=1110)
L2/3 IT_3 (n=460)
L2/3 IT_5 (n=374)
L2/3 IT_12 (n=349)
L2/3 IT_7 (n=93)
L2/3 IT_8 (n=64)
D
Comparing APP expression throughout disease progression in L2 IT Supertypes (increase) and L3 IT Supertypes (decrease)
Before Onset
Middle
Late
L2/3 IT_1
L2/3 IT_6
L2/3 IT_10
L2/3 IT_13
log1p Cp10k
Δ log1p Cp10k

Figure S5, related to Figure 6: Validation of spatial coordinate transfer and layer-specific gene expression across disease progression. A) Confusion matrix comparing each snRNA-seq cell's annotated Subclass to the Subclass of its nearest MERFISH neighbor in the PIANO latent space. Strong diagonal enrichment confirms accurate spatial coordinate assignment. B) Generated *MME* expression in vulnerable and unaffected SST Supertypes plotted separately across CPS, showing a decrease in upper cortical layers in both groups, with panels showing the difference relative to baseline (Early at CPS 0.2, Middle 0.5, and Late 1.0). C) Spatial distribution of L2/3 IT Supertypes across the representative MERFISH section, demonstrating that individual Supertypes occupy distinct cortical depths. D) Generated *APP* expression across CPS for L2-specific (L2/3 IT_1, L2/3 IT_6) and L3-specific (L2/3 IT_10, L2/3 IT_13) Supertypes, showing increased expression in L2 and decreased expression in L3.

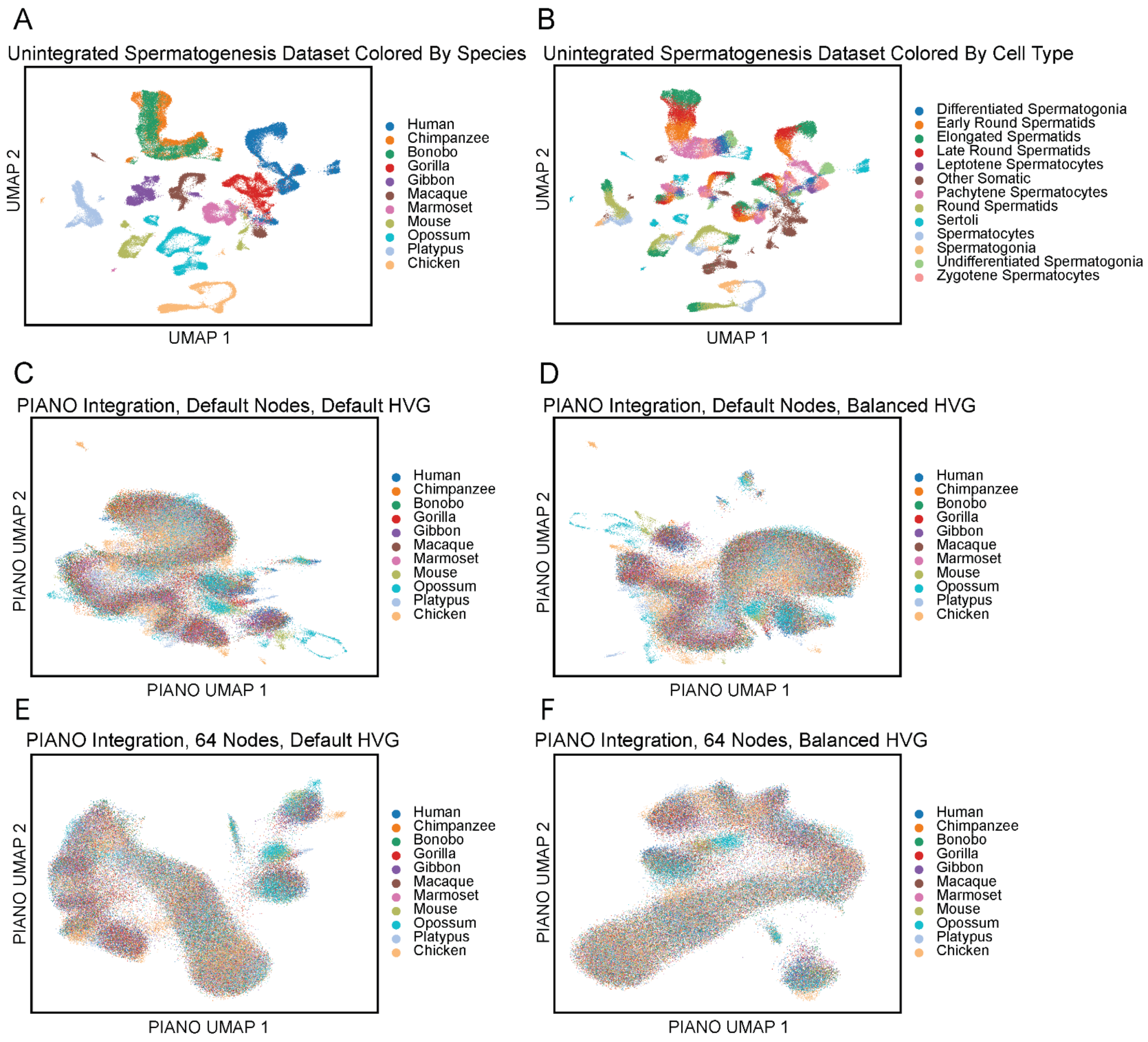


Figure S6, related to Figure 4: Spermatogenesis integration across evolutionarily distant species. A) The two dimensional UMAP projection of the PCA of the original counts data shows separation across species. B) The same UMAP shows evolutionary conservation of cell types, which are separated by species using PCA. C-D) A balanced (across evolutionarily clades) highly variable genes selection approach may lead to improvements over default highly variable gene selection (which would consider only species). E-F) A narrower model architecture may improve performance for small datasets with diverse species, regardless of gene selection.
