## Supplementary material for "PIANO: Probabilistic Inference Autoencoder Networks for multi-Omics enables robust generative modeling of gene expression and scales single-cell integration to 100 million cells": Key Resources Table.

| REAGENT or RESOURCE | SOURCE | IDENTIFIER |
|---|---|---|
| Deposited data | | |
| Pancreas data | Grün et al.[12]<br>Muraro et al.[13]<br>Lawlor et al.[14]<br>Segerstolpe et al.[15] | https://figshare.com/articles/dataset/Benchmarking_atlas-level_data_integration_in_single-cell_genomics_-_integration_task_datasets_Immune_and_pancreas_/12420968?file=24539828 |
| Lung data | Vieira Braga et al.[11] | https://figshare.com/articles/dataset/Benchmarking_atlas-level_data_integration_in_single-cell_genomics_-_integration_task_datasets_Immune_and_pancreas_/12420968?file=24539828 |
| Immune data | Oetjen et al.[17]<br>Datasets - Single Cell Gene Expression - Official 10x Genomics Support[18]<br>Freytag et al.[19]<br>Sun et al.[20]<br>Villani et al.[21] | https://figshare.com/articles/dataset/Benchmarking_atlas-level_data_integration_in_single-cell_genomics_-_integration_task_datasets_Immune_and_pancreas_/12420968?file=24539828 |
| Tabula muris data | Tabula Muris Consortium[22] | https://figshare.com/articles/dataset/Single-cell_RNA-seq_data_from_Smart-seq2_sequencing_of_FACS_sorted_cells_v2_/5829687/8 |
| Heart data | Kanemaru et al.[16] | https://figshare.com/articles/dataset/Batch_Alignment_of_single-cell_transcriptomics_data_using_Deep_Metric_Learning/20499630/2 |
| Primate Basal Ganglia data | Johansen, Fu et al.[29]<br>Dan, Turner et al.[30] | https://alleninstitute.github.io/HMBA_BasalGanglia_Consensus_Taxonomy/#rna-seq-data |
| Ortholog gene tables | Dyer et al.[62] | https://mart.ensembl.org/index.html |
| UniProtKB gene lists | Ahmad et al.[48] | https://www.uniprot.org/uniprotkb |
| AnimalTFDB4 gene lists | Shen et al.[49] | https://guolab.wchscu.cn/AnimalTFDB4/#/Download |
| Developmental mouse data | Schroeder et al.[32] | https://singlecell.broadinstitute.org/single_cell/study/SCP2719/a-multi-region-transcriptomic-atlas-of-developmental-cell-type-diversity-in-mouse-brain |
| Developmental marmoset data | Schroeder et al.[32] | https://singlecell.broadinstitute.org/single_cell/study/SCP2706/a-multi-region-transcriptomic-atlas-of-developmental-cell-type-diversity-in-marmoset-brain |
| Spermatogenesis data | Guo, Z.-H. et al.[35] | https://drive.google.com/drive/folders/1bNjY7x0ouzEWQhBFQZjKTFbPi6kSHr4L |
| Allen Institute whole mouse brain atlas single | Yao et al.[2] | https://alleninstitute.github.io/abc_atlas_access/descriptions/WMB_dataset.html |

| | | |
|---|---|---|
| cell RNA sequencing data | | |
| Broad Institute whole mouse brain MERFISH data | Zhang et al.[39] | https://alleninstitute.github.io/abc_atlas_access/descriptions/Zhuang_dataset.html |
| SEA-AD data | Gabbito et al.[41] | https://registry.opendata.aws/allen-sea-ad-atlas |
| Tahoe-100M | Zhang et al.[46] | https://www.biorxiv.org/content/10.1101/2025.02.20.639398v3.full |
| Software and algorithms | | |
| ScanPy 1.11.4 | Wolf et al.[64] | https://scanpy.readthedocs.io/en/stable/installation.html |
| PIANO | Current Manuscript | https://github.com/NingWang1729/piano https://pypi.org/project/piano-integration |
| Seurat 5.3.0 (Seurat V3 CCA) | Satija et al.[23] Butler et al.[8] Stuart et al.[40] | https://satijalab.org/seurat |
| Harmony 0.0.10 | Korsunsky et al.[7] | https://portals.broadinstitute.org/harmony |
| scIB benchmarking metrics | Luecken et al.[27] | https://scib-metrics.readthedocs.io/en/stable/#installation |
| scvi-tools v1.3.3 (scVI, sysVI) | Lopez et al.[24] Hrovatin et al.[25] | https://docs.scvi-tools.org/en/stable/installation.html |
| scDREAMER 0.0.0rc1 | Shree et al.[26] | https://github.com/Zafar-Lab/scDREAMER |
